# Specification dependence of EEG theta/alpha ratio associations with cognitive impairment: a multiverse analysis

**DOI:** 10.64898/2026.09.03.749187

**Authors:** Jinwon Chang, Seo Ho Song

**Affiliations:** Williams College, 39 Chapin Hall Drive, Williamstown, Massachusetts 01267, United States; Department of Psychiatry, Beth Israel Deaconess Medical Center, Harvard Medical School, Boston, Massachusetts 02215, United States

**Keywords:** electroencephalography, theta/alpha ratio, multiverse analysis, dementia, Parkinson’s disease, Alzheimer’s disease

## Abstract

**Objective:** The theta/alpha ratio, an index of electroencephalographic slowing, has been proposed as a marker of cognitive impairment, but estimates may depend on preprocessing, spectral, spatial, and statistical decisions. We used multiverse analysis to evaluate its robustness across plausible pipelines.

**Methods:** Two resting-state EEG datasets were analyzed. Dataset 1 included 49 controls and 100 participants with Parkinson’s disease spanning normal cognition to dementia. Dataset 2 included 29 controls, 36 patients with Alzheimer’s disease, and 23 with frontotemporal dementia. Eight decisions were varied: filtering, artifact correction, independent component analysis (ICA), referencing, spatial summary, alpha-band definition, spectral method, and covariate adjustment, yielding 1,280 specifications (universes).

**Results:** Findings varied across the unrestricted multiverse. In Dataset 1, all universes in the exploratory family combining ICA with absolute fast Fourier transform (FFT) or Welch power yielded p < 0.05 for comparisons of controls and cognitively normal Parkinson’s disease with Parkinson’s disease dementia and produced significant associations with Montreal Cognitive Assessment scores in more than 95% of universes. In Dataset 2, absolute FFT or Welch power supported control versus Alzheimer’s disease differences in more than 95% of universes, while ICA with absolute power yielded significant Mini-Mental State Examination associations in all specifications. Higher theta/alpha ratios were consistently associated with greater cognitive impairment.

**Conclusions:** The theta/alpha ratio is a promising but specification-dependent marker of cognitive impairment, contingent particularly on ICA and spectral power quantification.

**Significance:** Identifying the analytic conditions under which theta/alpha findings remain stable supports standardized, reproducible evaluation of quantitative EEG markers of cognitive impairment.

**Highlights:**

- Theta/alpha ratio findings varied substantially across 1,280 plausible EEG analysis pipelines.
- Independent component analysis with absolute spectral power yielded the most stable effects.
- Higher theta/alpha ratios consistently indicated greater cognitive impairment in robust pipelines.

## 1. Introduction

Quantitative electroencephalography (qEEG) has been widely investigated as a potential source of biomarkers for neurological and psychiatric disorders because it provides a noninvasive and accessible measure of brain activity. Quantitative measures have also been investigated in Alzheimer’s disease, frontotemporal dementia, Parkinson’s disease dementia, depression, and other clinical populations. However, translating qEEG findings into reliable clinical biomarkers remains difficult. Clinical samples vary in diagnosis, symptom severity, medication status, cognitive profile, and demographic characteristics, while EEG measures are additionally influenced by recording hardware, electrode montage, preprocessing, referencing, artifact correction, and feature-extraction methods. These combined sources of heterogeneity can substantially affect the reproducibility and clinical interpretation of qEEG findings (Babiloni et al., 2021).

Methodological variability is particularly important because a single EEG dataset can be processed using many defensible analytical pipelines. Filtering parameters, artifact-removal procedures, normalization, referencing, spatial selection, spectral estimation, and statistical adjustment may each alter the resulting effect. Although standardized or optimal EEG-processing procedures have been proposed, many methodological recommendations were developed primarily using healthy or nonclinical samples, leaving uncertainty regarding whether the same procedures are optimal in populations with altered neural physiology (Delorme & Makeig, 2004). Consequently, disagreement between clinical EEG studies may reflect genuine biological heterogeneity, methodological differences, or an interaction between the two.

This problem forms part of the broader challenge of scientific replicability. The replication crisis has been widely debated across the social, behavioral, and biomedical sciences (Romero, 2019). This concern is particularly relevant to clinical research, where abnormalities may be small, heterogeneous, and susceptible to methodological artifacts (Clayson, 2024). Neurophysiological and neuroimaging analyses are especially vulnerable because EEG and functional magnetic resonance imaging require numerous processing and modeling decisions. Indeed, when independent research teams analyzed the same neuroimaging dataset, differences in analytical workflows produced substantial variability in inferential results (Botvinik-Nezer et al., 2020). Meta-analysis can estimate the overall direction of effects across studies, but it cannot fully determine whether an observed finding depends on the raw-data processing and statistical decisions made within the contributing studies.

Multiverse analysis provides a direct framework for examining this analytical uncertainty (Clayson, 2024; Kołodziej et al., 2021). In this framework, a universe represents one complete and internally consistent analytical specification comprising a particular combination of preprocessing choices, parameter settings, feature-extraction procedures, and statistical models. The multiverse consists of the full set of universes generated from prespecified combinations of defensible analytical decisions. Whereas a conventional analysis reports the result obtained from one selected pipeline, multiverse analysis evaluates how effect estimates and inferential outcomes vary across alternative specifications. This allows investigators to determine whether a finding is robust across reasonable analytical choices or depends on a restricted subset of methodological decisions.

Related guided multiverse approaches embed analytical pipelines in a lower-dimensional space and use active learning to identify informative or high-performing specifications without evaluating every possible pipeline (Dafflon et al., 2022). In the present study, however, the number of candidate pipelines was computationally manageable. We therefore used an exhaustive multiverse, allowing all prespecified combinations to be evaluated directly. This design permits the distribution of results to be stratified by individual analytical decisions and their combinations without relying on model-based estimates of unsampled universes.

The theta/alpha ratio provides a useful clinical EEG measure for evaluating such analytical dependence. It has been proposed as an index of EEG slowing, reflecting a relative shift from higher-frequency alpha activity toward lower-frequency theta activity (Hamilton et al., 2021; Liang et al., 2026). Elevated theta/alpha ratios have been investigated in disorders associated with cognitive impairment, including Alzheimer’s disease and frontotemporal dementia (Fahimi et al., 2017; Schmidt et al., 2013). However, the ratio is not a methodologically invariant quantity. Its estimated magnitude may be influenced by the spectral algorithm, use of absolute or relative normalization, alpha-band definition, EEG reference, and artifact-removal strategy (Chang, 2025; Yao et al., 2005). Consequently, an apparently significant clinical association obtained using one pipeline may not necessarily persist under alternative, equally defensible analytical specifications.

The present study therefore applied an exhaustive multiverse framework to examine the robustness of the theta/alpha ratio as a marker of cognitive impairment across two independent clinical EEG datasets. The first dataset included healthy controls and patients with Parkinson’s disease spanning cognitively normal, mild cognitive impairment, and dementia subgroups, whereas the second included healthy controls and patients with Alzheimer’s disease or frontotemporal dementia. Across both datasets, the multiverse systematically varied filtering, artifact removal, independent component analysis, referencing, spatial aggregation, alpha-band definition, spectral estimation, and covariate adjustment. We examined whether group differences and cognitive associations remained consistent across plausible analytical universes, which analytical decisions most strongly influenced the resulting inferences, and whether findings meeting the ≥95% robustness criterion also showed consistent directions of Cohen’s *d* or standardized regression coefficients.

## 2. Methods

### 2.1. Dataset selection

Publicly available EEG datasets were selected using prespecified criteria to permit a common multiverse analysis across clinically relevant populations with cognitive impairment. PubMed and IEEE Xplore were searched using terms related to dementia, Alzheimer’s disease, mild cognitive impairment, frontotemporal dementia, Lewy body dementia, EEG/qEEG, and biomarker or diagnostic applications.

Eligible datasets required participant-level resting-state scalp EEG, defined diagnostic or cognitive-status labels, a cognitively healthy comparison group, cognitive measures such as the Montreal Cognitive Assessment (MoCA) or Mini-Mental State Examination (MMSE), demographic variables including age and sex, and sufficient recording information to implement the same core processing decisions. Datasets were excluded when only derived EEG features were available, clinical or demographic information was insufficient, only task-related EEG was provided, or the processing pipeline could not be reconstructed.

Two datasets met these criteria. Dataset 1 included healthy controls and patients with Parkinson’s disease spanning cognitively normal, mild cognitive impairment, and dementia groups with MoCA scores. Dataset 2 included healthy controls and patients with Alzheimer’s disease or frontotemporal dementia with MMSE scores. The datasets were considered complementary rather than direct replications because they differed in population, recording duration, eye state, electrode density, and cognitive assessment.

The present study was a secondary analysis of de-identified, publicly available EEG datasets and did not involve new participant recruitment or data collection; the analysis was determined to be exempt from additional institutional review at the authors’ institutions.

### 2.2. Multiverse design

EEG results can depend on method- and parameter-level choices, whose effects may vary with sampling rate, recording duration, electrode density, montage, and data quality (Robbins et al., 2020; Huang et al., 2025; Wiesman et al., 2022; Winkler et al., 2015). The multiverse therefore focused on major method-level decisions that could be applied consistently across both datasets, while parameters such as spectral-window duration and artifact thresholds were held constant where possible.

The evaluated decisions were filter type, Artifact Subspace Reconstruction (ASR), independent component analysis (ICA), reference, spatial summary, alpha-band definition, spectral estimation and normalization, and demographic covariate adjustment. The aim was not to identify a universally optimal pipeline, but to determine whether theta/alpha findings remained stable across defensible analytical choices.

### 2.3. Dataset 1

#### 2.3.1. Participants and acquisition

Dataset 1 comprised 100 patients with Parkinson’s disease (PD) and 49 healthy controls (Singh et al., 2023) (Table 1). Patients were evaluated by a movement-disorders specialist and met United Kingdom Parkinson’s Disease Society Brain Bank criteria (Gibb & Lees, 1988).

**Table 1.** Clinical demographic characteristics of Parkinson’s disease subgroups and healthy controls.

|  | Control | PD-CN | PDD | PDMCI | P value |
| --- | --- | --- | --- | --- | --- |
| N | 49 | 47 | 19 | 34 |  |
| Sex (% female) | 46.9 | 42.6 | 15.8 | 26.5 | 0.046 |
| Age (years) | 70.9 ± 1.1 | 66.0 ± 1.1 | 70.2 ± 1.9 | 71.2 ± 1.4 | 0.005 |
| MoCA | 26.7 ± 0.3 | 27.4 ± 0.2 | 17.8 ± 0.8 | 23.7 ± 0.2 | <0.001 |
| UPDRS | — | 11.2 ± 1.0 | 11.9 ± 1.1 | 16.5 ± 1.9 | 0.044 |
| LEDD | — | 758.8 ± 54.9 | 1065.8 ± 130.5 | 750.3 ± 75.3 | 0.024 |
| Disease Duration | — | 4.8 ± 0.6 | 4.8 ± 1.0 | 4.4 ± 0.5 | 0.883 |

Data are expressed as mean ± standard error of mean. Sex was expressed as the proportion of female participants. MoCA, Montreal Cognitive Assessment; UPDRS, motor Unified Parkinson’s Disease Rating Scale; LEDD, Levodopa Equivalent Daily Dose; PD-CN, cognitively normal Parkinson’s disease; PDD, Parkinson’s disease dementia; PDMCI, Parkinson’s disease with mild cognitive impairment.

Written informed consent was obtained under procedures approved by the University of Iowa Human Subjects Review Board (IRB #201707828).

Patients were assessed in the ON-medication state, and levodopa-equivalent daily dose (LEDD) was available. Cognitive status was assessed using the MoCA (Dalrymple-Alford et al., 2010). Parkinson’s disease dementia (PDD) was defined as MoCA <22, Parkinson’s disease with mild cognitive impairment (PDMCI) as MoCA 22–25, and cognitively normal Parkinson’s disease (PD-CN) as MoCA ≥26. Patients also completed the motor Unified Parkinson’s Disease Rating Scale (UPDRS-III) (Goetz et al., 2008).

EEG data were obtained from OpenNeuro (doi:10.18112/openneuro.ds004584.v1.0.0). Approximately 2 min of eyes-open resting EEG were recorded using a 64-channel actiCAP system at 500 Hz, with Pz as reference and Fpz as ground. Data were imported into EEGLAB and resampled to 250 Hz before multiverse processing.

### 2.4. EEG preprocessing decision space

#### 2.4.1. Filtering and ASR

A 1-Hz high-pass filter was implemented using either finite impulse response (FIR) or Butterworth filtering. High-pass filtering can improve ICA decomposition by reducing slow drift and low-frequency nonstationarity (Klug & Gramann, 2020; Winkler et al., 2015). FIR and Butterworth filters were compared because they involve different trade-offs in phase preservation, attenuation, filter order, and edge effects (Widmann et al., 2015). Because the analysis focused on band-limited spectral power rather than phase-sensitive measures, both were treated as defensible alternatives.

ASR was either applied or omitted. ASR identifies high-variance signal subspaces that deviate from a reference covariance structure and reconstructs contaminated data (Miyakoshi, 2023). When applied, burst artifacts were detected using a 0.5-s sliding window and a 20-SD threshold, with bad segments rejected using root-mean-square thresholds of −Inf to 7 and a maximum of 25% channel outliers. The 20-SD criterion is consistent with previous ASR evaluations (Chang et al., 2019). Recording durations exceeded the approximately 40 s reported as sufficient for reliable estimation of many qEEG features (Gudmundsson et al., 2007).

#### 2.4.2. Independent component analysis and reference

Extended infomax ICA was either applied or omitted using EEGLAB runica. ICA is widely used to separate ocular, muscular, cardiac, and other non-neural sources (Vigário et al., 2000; Grech et al., 2008). Components classified by ICLabel as ocular, cardiac, line-noise, or muscle-related with probability ≥80% were removed. Because ICA may attenuate artifacts but can also remove physiologically meaningful activity, it was treated as an analytical branch rather than an assumed optimal procedure (Grech et al., 2008).

EEG was then re-referenced using either average reference or the reference electrode standardization technique (REST) (Yao, 2001). Average reference depends on sufficiently broad and approximately uniform scalp coverage, whereas REST estimates potentials relative to a theoretical point at infinity using a head-model transformation (Yao et al., 2019; Qin et al., 2010). The comparison therefore tested whether theta/alpha findings were sensitive to reference choice.

### 2.5. Spectral decision space

#### 2.5.1. Frequency bands and spectral estimation

Theta power was defined as 4–8 Hz, while alpha was defined as either 8–12 or 8–13 Hz. The clinical EEG glossary broadly defines alpha within 8–13 Hz (Kane et al., 2017), although narrower definitions and alpha sub-bands are also commonly used (İşoğlu-Alkaç & Strüber, 2006). Fixed boundaries may be less appropriate in neurodegenerative disorders because the dominant rhythm can slow, but individualized alpha peaks may also be difficult to identify consistently and may not reliably distinguish Alzheimer’s disease from healthy aging (Kopčanová et al., 2024). We therefore compared two conventional fixed definitions.

Spectral analyses used 2-s windows with 50% overlap. Power was estimated using either direct FFT-based analysis or Welch’s method. Welch estimates used Hamming windows without detrending. Although both rely on Fourier decomposition, Welch averaging reduces variance and sensitivity to transient fluctuations, potentially yielding different theta/alpha estimates.

#### 2.5.2. Absolute, relative-normalized, and aperiodic-adjusted power

Five spectral methods were evaluated: absolute Welch power, relative-normalized Welch power, absolute FFT power, relative-normalized FFT power, and aperiodic-adjusted power estimated using Fitting Oscillations and One-Over-F (FOOOF) (Donoghue et al., 2020).

For absolute-power branches, the theta/alpha ratio was calculated as theta-band power divided by alpha-band power. For relative-normalized branches, this ratio was subsequently divided by total spectral power from 1–80 Hz. Thus, relative normalization refers to normalization of the theta/alpha ratio itself rather than division of separately normalized theta and alpha bands. This approach follows previous work comparing absolute and relative spectral power ratios (Chang, 2025). Test-retest analyses have also shown that spectral ratios can be highly reliable and that absolute and relative formulations may differ in reliability, supporting their treatment as distinct qEEG features (Chang, 2025).

FOOOF was included because conventional spectra contain both periodic oscillations and broadband aperiodic activity. It was fit from 1–40 Hz using a fixed aperiodic model, peak-width limits of 1–8 Hz, a maximum of six peaks, minimum peak height of 0.05, and a peak threshold of 2 (Donoghue et al., 2020). Comparing absolute, relative-normalized, and aperiodic-adjusted measures therefore tested whether clinical associations depended on global spectral magnitude or the aperiodic background.

### 2.6. Spatial decision space

Four spatial summaries were evaluated: Fz, Cz, a frontal region of interest comprising Fz, F3, F4, F7, F8, Fp1, and Fp2, and a central region comprising Cz, C3, and C4.

Regional averages can reduce channel-specific noise, whereas single-electrode measures may be useful for low-density systems. Frontal and central regions were selected because frontal theta activity is associated with working memory and cognitive control (Jensen & Tesche, 2002; Cavanagh & Frank, 2014), while frontal alpha has been studied in relation to cognition, emotion, and psychopathology (Allen et al., 2018). In Parkinson’s disease, mid-frontal theta abnormalities have also been reported and may be relatively insensitive to levodopa in cognitive-control paradigms (Singh et al., 2018).

### 2.7. Statistical analysis and universe construction

For Dataset 1, clinical and demographic characteristics were compared across CTL, PD-CN, PDMCI, and PDD using ANOVA with Tukey-adjusted comparisons for age, disease duration, and LEDD; Kruskal–Wallis tests with Holm-adjusted comparisons for MoCA and UPDRS-III; and chi-square tests for sex.

Within each universe, theta/alpha group differences were evaluated using ANOVA or ANCOVA adjusting for age and sex, followed by Tukey-adjusted pairwise comparisons.

Associations with MoCA and UPDRS-III were evaluated using linear regression with and without demographic adjustment. Because PD cognitive groups were defined using MoCA thresholds, the MoCA regression was treated as a dimensional consistency analysis rather than independent validation.

The full decision space included filter type (2 levels), ASR (2), ICA (2), reference (2), spatial summary (4), alpha definition (2), spectral method (5), and covariate adjustment (2), yielding 1,280 universes per outcome.

### 2.8. Dataset 2

#### 2.8.1. Participants, acquisition, and analysis

Dataset 2 included 36 patients with Alzheimer’s disease (AD), 23 with frontotemporal dementia (FTD), and 29 healthy controls (CTL) (Miltiadous et al., 2023) (Table 2). Cognitive function was assessed using the MMSE (Kurlowicz & Wallace, 1999). Diagnoses were based on DSM-III-R, DSM-IV, ICD-10, and NINCDS–ADRDA criteria (Bell, 1994; McKhann et al., 1984). The original study obtained informed consent and was approved by the Scientific and Ethics Committee of AHEPA University Hospital, Aristotle University of Thessaloniki (protocol 142/12-04-2023).

**Table 2.** Clinical demographic characteristics of dementia patients and healthy controls.

|  | CTL | FTD | AD | P value |
| --- | --- | --- | --- | --- |
| N | 29 | 23 | 36 |  |
| Sex (% female) | 37.9 | 39.1 | 66.6 | 0.034 |
| Age (years) | 67.90 ± 1.00 | 63.65 ± 1.71 | 66.39 ± 1.32 | 0.115 |
| MMSE | 30.00 ± 0.00 | 22.17 ± 0.54 | 17.75 ± 0.75 | <0.001 |

Data are expressed as mean ± standard error of mean. Sex was expressed as the proportion of female participants. MMSE, Mini-Mental State Examination; CTL, healthy controls; AD, Alzheimer’s disease; FTD, frontotemporal dementia

Data were obtained from OpenNeuro (doi:10.18112/openneuro.ds004504.v1.0.9). Eyes-closed resting EEG was recorded using a Nihon Kohden 2100 system with 19 scalp electrodes at 500 Hz. Recording duration averaged approximately 13.5 min for AD, 12.0 min for FTD, and 13.8 min for CTL. These differences from Dataset 1 in eye state, duration, and electrode density were properties of the original datasets rather than multiverse branches.

The same preprocessing, spectral, spatial, and statistical decision structure was applied to Dataset 2. Age was evaluated using ANOVA with Tukey-adjusted comparisons, MMSE using Kruskal–Wallis tests with Holm-adjusted comparisons, and sex using chi-square tests. Within each universe, group differences were tested using ANOVA or ANCOVA with age and sex, and MMSE associations using linear regression with and without demographic adjustment.

REST findings were interpreted cautiously because the 19-channel montage provides sparser spatial sampling than Dataset 1, potentially affecting both average-reference and head-model-based transformations (Yao et al., 2019; Qin et al., 2010).

### 2.9. Robustness criteria

Results were summarized across universes rather than from a single preferred pipeline. The primary descriptive measure was the proportion of universes yielding *p* < 0.05, with ≥95% treated as high stability within the evaluated decision space. This threshold was descriptive rather than a formal probability of a true effect.

Robustness was also evaluated using Cohen’s *d* for group contrasts and standardized regression coefficients for cognitive associations, with emphasis on consistency of effect direction. Because universes were derived from the same participants, they were statistically dependent and were not treated as independent hypothesis tests. Pipeline families identified after inspection of the multiverse, particularly ICA combined with absolute FFT or Welch power, were considered exploratory and require independent prospective validation.

## 3. Results

### 3.1. Dataset 1

Age differed across groups (ANOVA, *F*(3,145) = 4.42, *p* = 0.005), with PD-CN participants younger than CTL and PDMCI participants after Tukey adjustment. Sex distribution also differed (χ²(3) = 7.99, *p* = 0.046). MoCA differed across groups (*H*(3) = 99.40, *p* < 0.001), as expected from the MoCA-based cognitive classification, and UPDRS-III differed among PD subgroups (*H*(2) = 6.24, *p* = 0.044) (Table 1).

Across all 1,280 universes, inferential results showed substantial specification dependence (Figure 1). Significant results occurred in 40% of universes for CTL versus PDD, 38% for PD-CN versus PDD, 17% for CTL versus PDMCI, 49% for MoCA, and 9% for UPDRS-III. The strongest results were concentrated in pipelines combining ICA with absolute spectral power (Figure 2). All ICA + absolute Welch universes and all ICA + absolute FFT universes were significant for CTL versus PDD; corresponding proportions for PD-CN versus PDD were 100% and 98%. MoCA associations were significant in 100% of both absolute-power families. Relative-power and FOOOF specifications produced substantially less consistent evidence.

**Figure 1.**
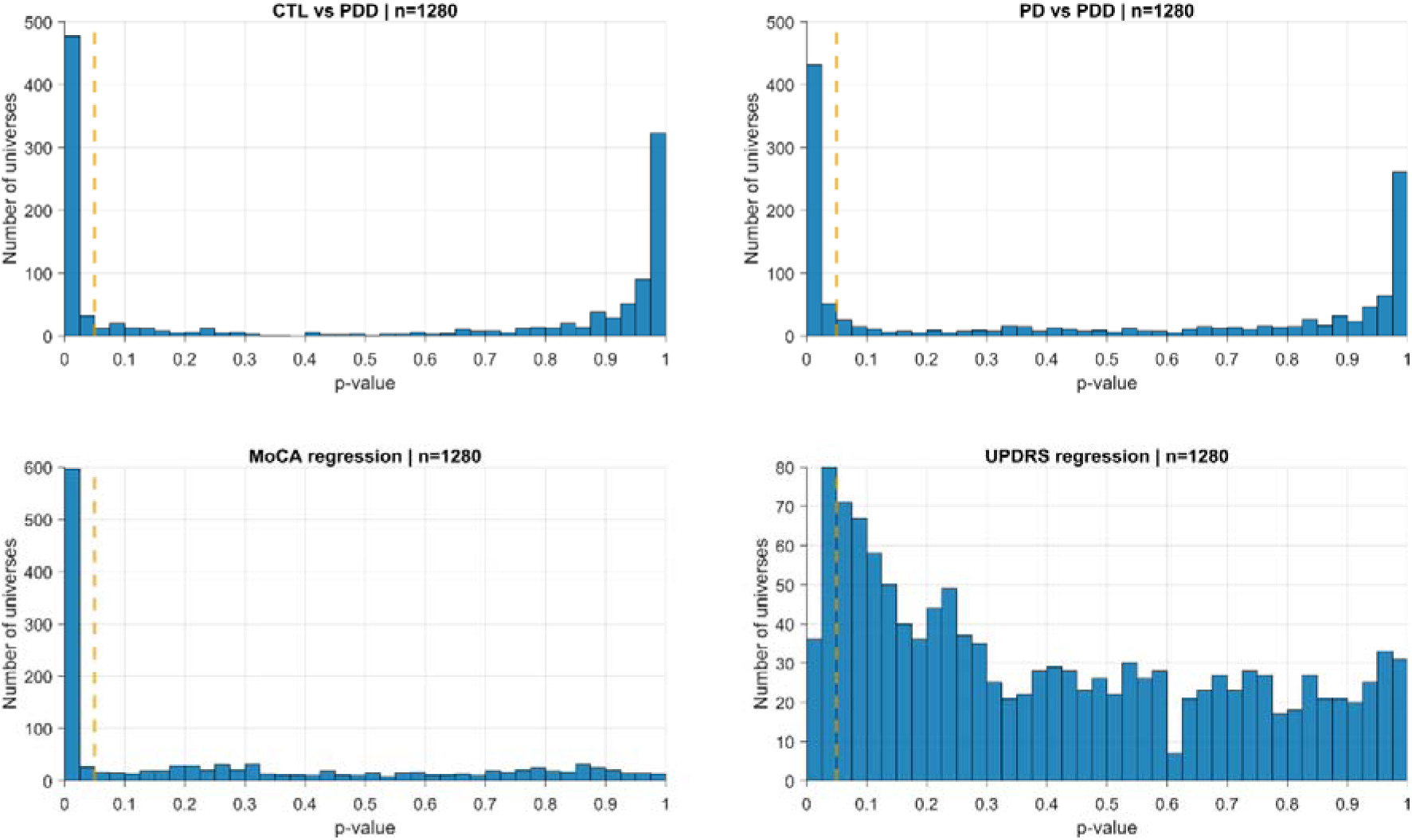
Dataset 1: p-value distributions across all 1,280 universes for CTL versus PDD, PD-CN versus PDD, MoCA regression, and UPDRS-III regression. The dashed line marks p = 0.05. PD in the figure denotes cognitively normal Parkinson’s disease (PD-CN). CTL, healthy controls; MoCA, Montreal Cognitive Assessment; PD-CN, cognitively normal Parkinson’s disease; PDD, Parkinson’s disease dementia; UPDRS-III, motor Unified Parkinson’s Disease Rating Scale.

**Figure 2.**
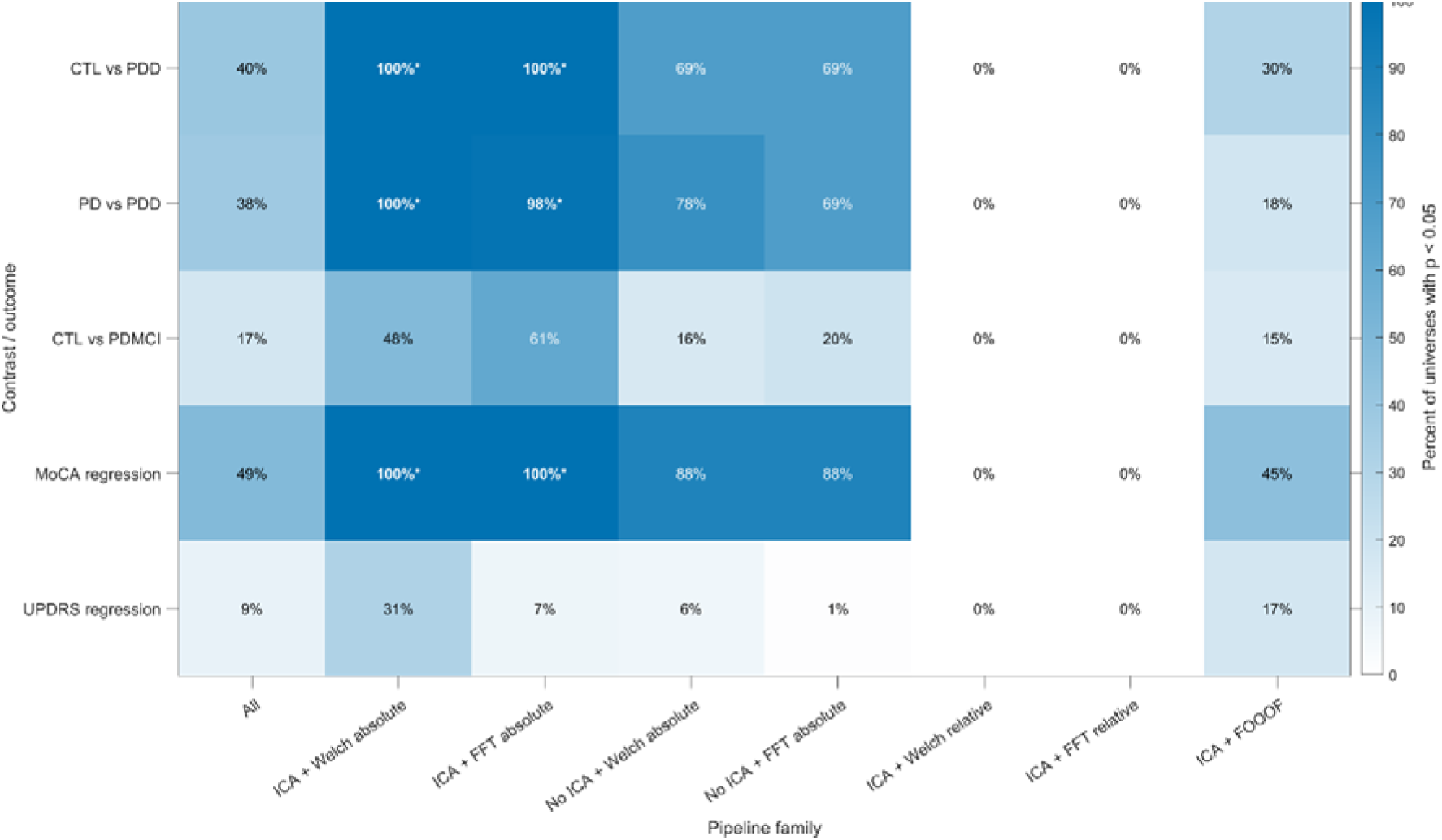
Dataset 1: percentage of universes with p < 0.05 across the full multiverse and selected pipeline families. Asterisks indicate pipeline-family/outcome combinations meeting the prespecified ≥95% robustness criterion. PD in the figure denotes cognitively normal Parkinson’s disease (PD-CN). CTL, healthy controls; FFT, fast Fourier transform; FOOOF, Fitting Oscillations and One-Over-F; ICA, independent component analysis; MoCA, Montreal Cognitive Assessment; PD-CN, cognitively normal Parkinson’s disease; PDD, Parkinson’s disease dementia; PDMCI, Parkinson’s disease with mild cognitive impairment; UPDRS-III, motor Unified Parkinson’s Disease Rating Scale.

Within the exploratory ICA + absolute Welch/FFT family, all 256 CTL-versus-PDD and PD-CN-versus-PDD Cohen’s *d* estimates were negative with 95% confidence intervals excluding zero, indicating higher theta/alpha ratios in PDD given the contrast direction (CTL minus PDD and PD-CN minus PDD). All 256 standardized MoCA coefficients were also negative, indicating higher ratios with poorer cognition (Figure 3). Additional pairwise and decision- stratified results are provided in Supplementary Figures 1–8.

**Figure 3.**
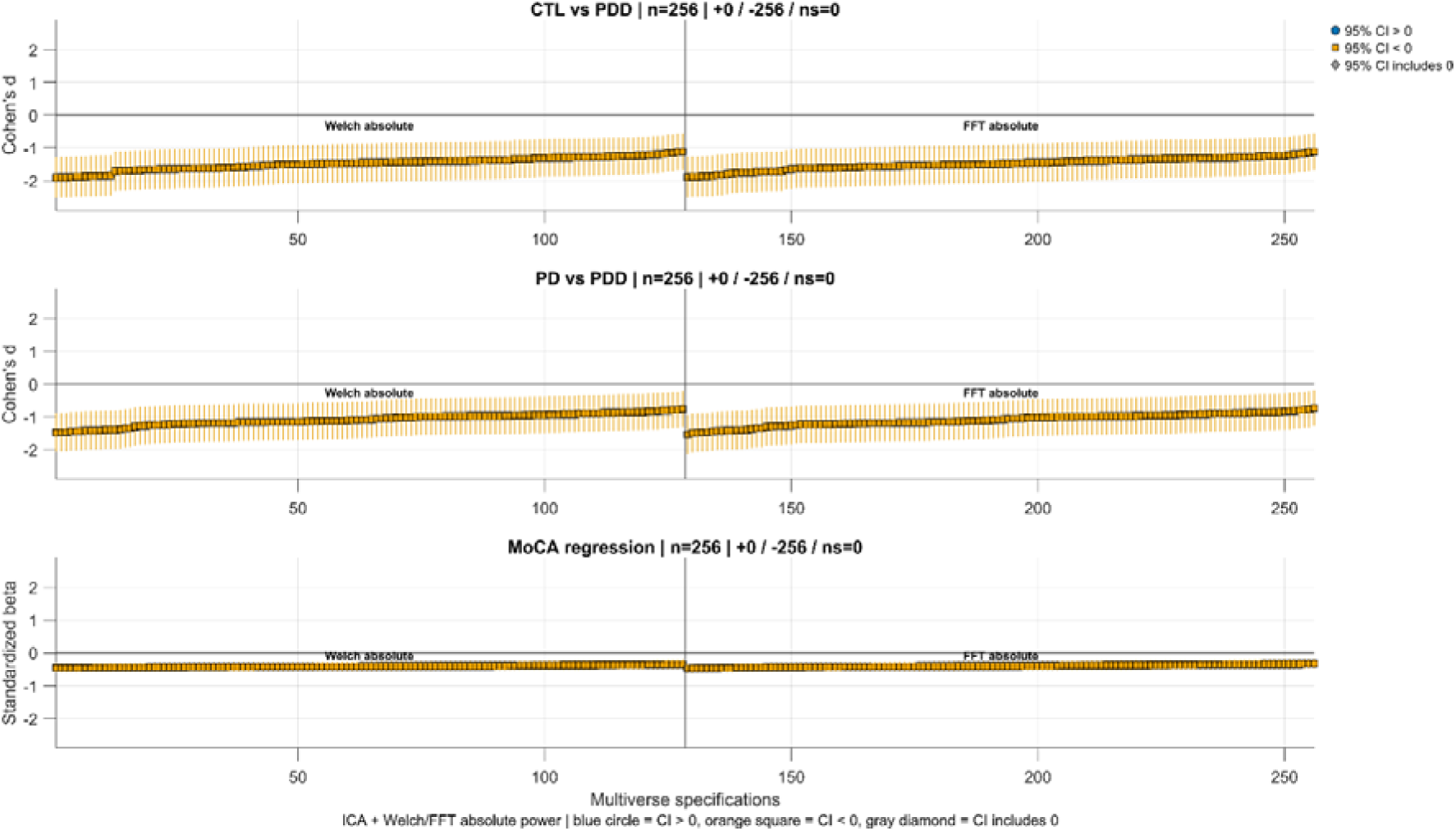
Dataset 1: specification curves for the exploratory ICA + absolute Welch/FFT family. Points show Cohen’s d for CTL versus PDD and PD-CN versus PDD, or standardized beta for MoCA regression; vertical bars show 95% confidence intervals. All 256 estimates in each panel were negative and their 95% confidence intervals excluded zero. CTL, healthy controls; FFT, fast Fourier transform; ICA, independent component analysis; MoCA, Montreal Cognitive Assessment; PD-CN, cognitively normal Parkinson’s disease; PDD, Parkinson’s disease dementia.

### 3.2. Dataset 2

Age did not differ across groups (*F*(2,85) = 2.22, *p* = 0.115), whereas sex distribution differed (χ²(2) = 6.78, *p* = 0.034). MMSE differed across groups (*H*(2) = 67.76, *p* < 0.001), with all Holm-adjusted pairwise comparisons significant (Table 2).

Across all universes, 67% yielded *p* < 0.05 for CTL versus AD, 32% for CTL versus FTD, 15% for AD versus FTD, and 79% for MMSE (Figures 4–5). As in Dataset 1, absolute-power families produced the most stable results. All ICA + absolute Welch and ICA + absolute FFT universes were significant for both CTL versus AD and MMSE. Relative-power families showed intermediate robustness, whereas FOOOF yielded substantially fewer significant universes.

**Figure 4.**
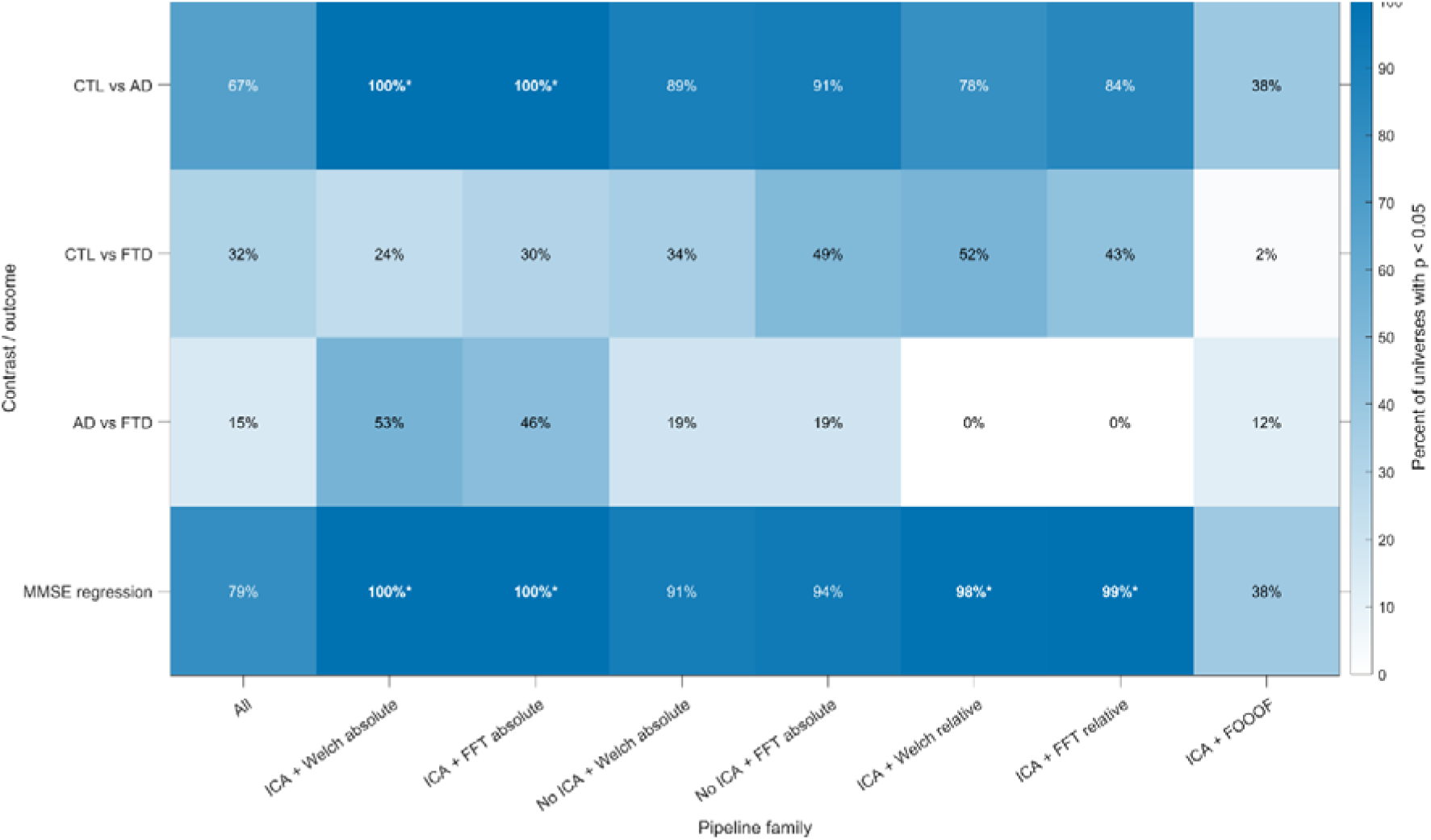
Dataset 2: percentage of universes with p < 0.05 across the full multiverse and selected pipeline families. Asterisks indicate combinations meeting the prespecified ≥95% robustness criterion. AD, Alzheimer’s disease; CTL, healthy controls; FFT, fast Fourier transform; FOOOF, Fitting Oscillations and One-Over-F; FTD, frontotemporal dementia; ICA, independent component analysis; MMSE, Mini-Mental State Examination.

**Figure 5.**
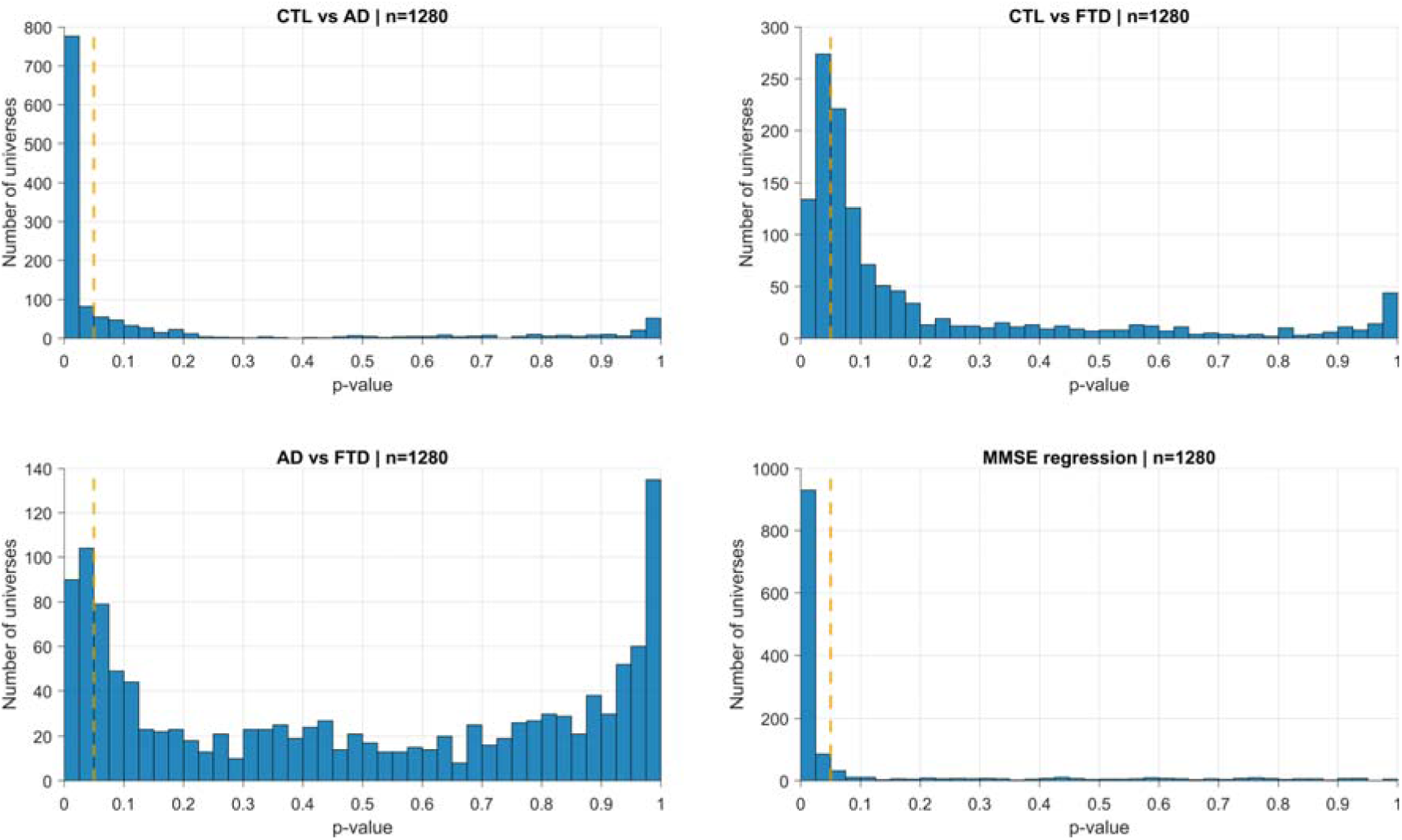
Dataset 2: p-value distributions across all 1,280 universes for CTL versus AD, CTL versus FTD, AD versus FTD, and MMSE regression. The dashed line marks p = 0.05. AD, Alzheimer’s disease; CTL, healthy controls; FTD, frontotemporal dementia; MMSE, Mini-Mental State Examination.

Directional consistency was also complete within the exploratory ICA + absolute Welch/FFT family: all 256 CTL-versus-AD Cohen’s *d* estimates and all 256 standardized MMSE coefficients were negative with confidence intervals excluding zero, indicating higher theta/alpha ratios in AD and with poorer cognitive performance (Figure 6). Additional group comparisons are presented in Supplementary Figures 9–11.

**Figure 6.**
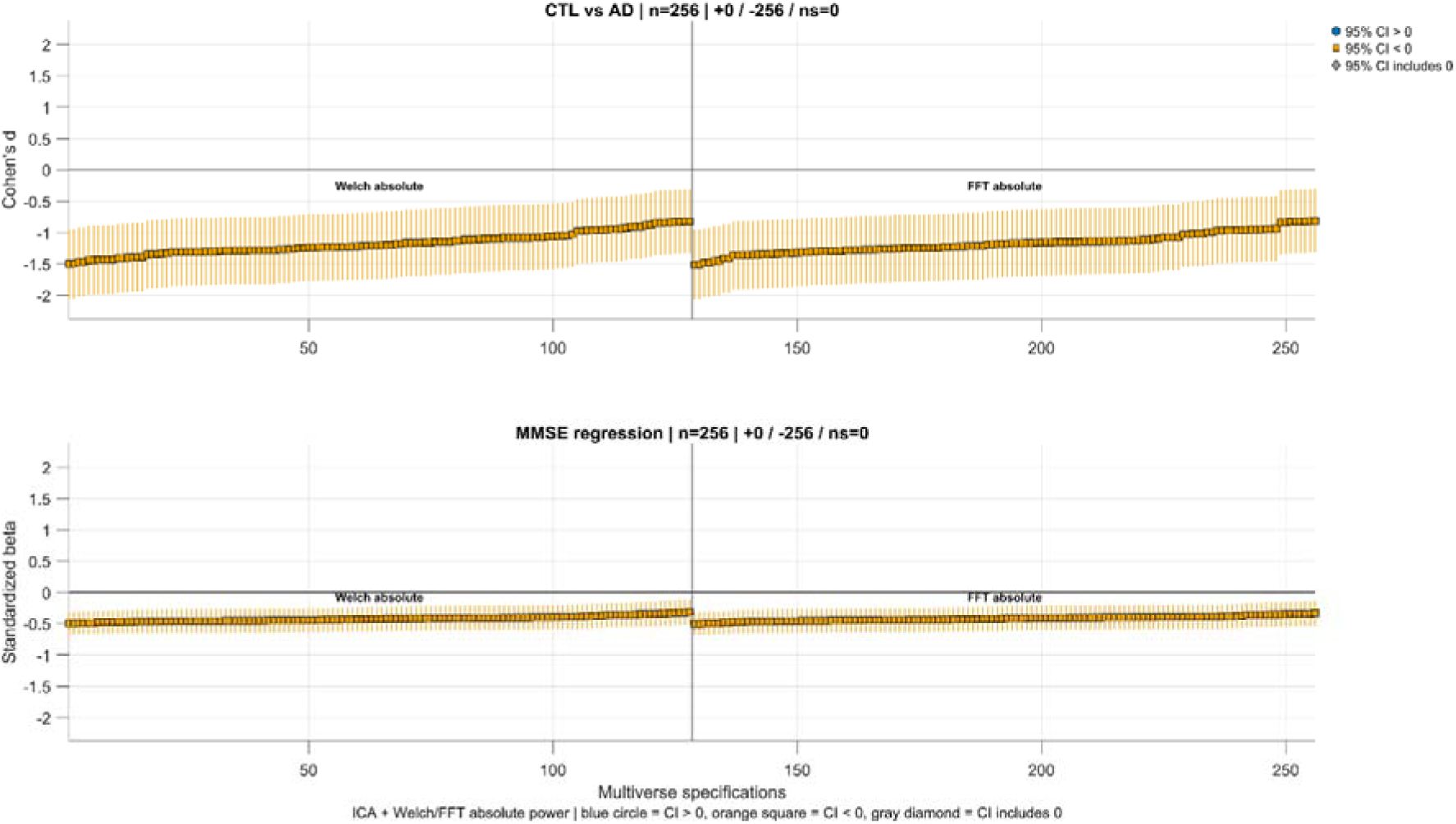
Dataset 2: specification curves for the exploratory ICA + absolute Welch/FFT family. Points show Cohen’s d for CTL versus AD or standardized beta for MMSE regression; vertical bars show 95% confidence intervals. All 256 estimates in each panel were negative and their 95% confidence intervals excluded zero. AD, Alzheimer’s disease; CTL, healthy controls; FFT, fast Fourier transform; ICA, independent component analysis; MMSE, Mini-Mental State Examination.

### 3.3. Decision-level sensitivity across datasets

Spectral quantification produced the largest decision-level differences across datasets (Figure 7). In Dataset 1, absolute Welch and FFT power yielded significant results in approximately 83–94% of universes for the principal PDD and MoCA outcomes, whereas relative power produced few or no significant universes and FOOOF showed intermediate performance. In Dataset 2, absolute Welch and FFT power yielded approximately 95–97% significant universes for CTL versus AD and MMSE. ICA further increased the proportion of significant universes, particularly in Dataset 2, whereas filter type, reference, spatial summary, and alpha definition had smaller effects. Across both datasets, conventional absolute spectral power was therefore the most consistent decision-level correlate of inferential robustness, with an additional contribution from ICA.

**Figure 7.**
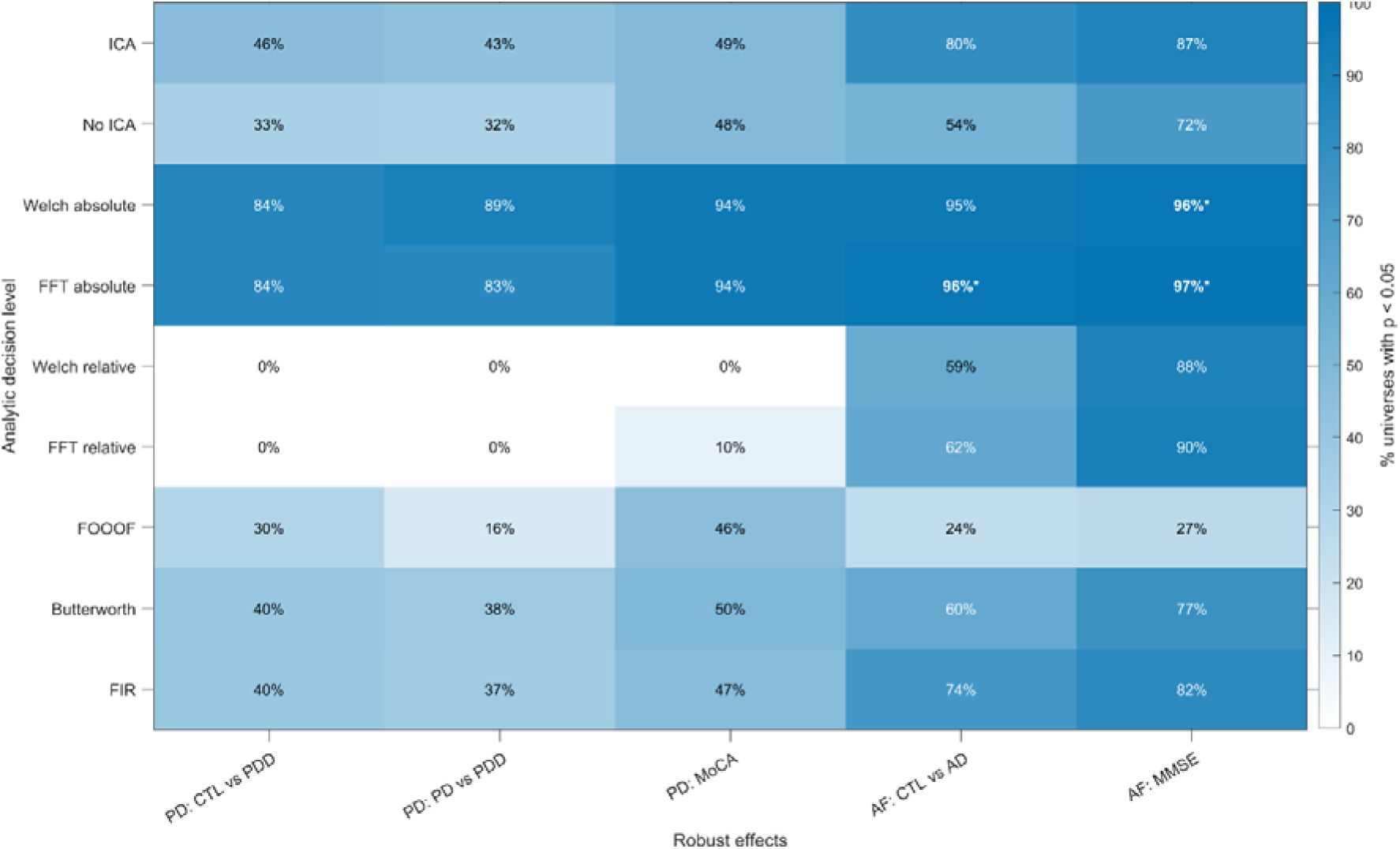
Percentage of universes with p < 0.05 stratified by selected analytic decision levels for the principal robust effects in Dataset 1 (Parkinson’s disease) and Dataset 2 (Alzheimer’s disease and frontotemporal dementia). Asterisks indicate decision-level subsets meeting the ≥95% robustness criterion. AD, Alzheimer’s disease; CTL, healthy controls; FFT, fast Fourier transform; FIR, finite impulse response; FOOOF, Fitting Oscillations and One-Over-F; ICA, independent component analysis; MMSE, Mini-Mental State Examination; MoCA, Montreal Cognitive Assessment; PD-CN, cognitively normal Parkinson’s disease; PDD, Parkinson’s disease dementia.

## 4. Discussion

### 4.1. Principal findings

The present study evaluated the theta/alpha ratio across 1,280 preprocessing, spectral, spatial, and statistical specifications in two clinical EEG datasets. Three findings were consistent.

First, pipeline families in which at least 95% of universes yielded *p* < 0.05 also showed stable effect directions, with higher theta/alpha ratios associated with greater cognitive impairment. Second, evidence was strongest for comparisons involving more advanced impairment, including CTL versus PDD, PD-CN versus PDD, and CTL versus AD. Third, ICA and spectral power estimation, particularly absolute FFT or Welch power, were most consistently associated with significant results, whereas filtering, reference, spatial summary, and alpha-band definition had smaller effects within the evaluated ranges.

These findings support the theta/alpha ratio as a promising marker of cognitive impairment, but also show that its robustness depends on analytical specification. The full multiverse did not support every group contrast or clinical association. Instead, stable evidence emerged within identifiable pipeline families. Thus, multiverse analysis does not identify one universally correct pipeline, but reveals whether findings are broadly robust or dependent on restricted methodological choices.

### 4.2. Theta/alpha ratio and cognitive impairment

EEG slowing is characteristic of several neurodegenerative disorders, and the theta/alpha ratio summarizes the relative shift from alpha toward theta activity. In Dataset 1, the most stable differences involved PDD rather than PDMCI or PD-CN, suggesting greater sensitivity to advanced cognitive impairment. Dataset 2 similarly showed robust CTL versus AD differences but not CTL versus FTD or AD versus FTD. This pattern is more consistent with global cognitive impairment than diagnostic specificity and is supported by the negative associations with MoCA and MMSE scores.

The datasets should nevertheless be viewed as complementary rather than direct replications. They differed in diagnosis, electrode density, recording system, cognitive assessment, and clinical context. Dataset 1 also included patients with PD assessed in the ON-medication state, whereas Dataset 2 included AD and FTD. Similar directional effects across these settings strengthen the relevance of EEG slowing but do not establish diagnostic specificity.

### 4.3. Preprocessing multiverse: sensitivity to ICA

Preprocessing decisions are important because EEG artifacts overlap spectrally and spatially with neural signals. Among the tested choices, ICA showed the clearest association with stable group and cognitive-score findings. ICA was developed for source separation and is widely used for artifact identification and neural-source estimation (Vigário et al., 2000; Delorme & Makeig, 2004). However, EEG source localization remains constrained by the inverse problem (Grech et al., 2008), and anatomical heterogeneity may further limit source-level interpretation in clinical samples. ICA is therefore often used primarily to identify ocular, muscular, cardiac, and other non-neural components (Romero et al., 2003).

In the present datasets, ICA-based artifact removal was associated with more consistent theta/alpha findings. This should not be interpreted as evidence that ICA is universally necessary. ICA can alter theta and alpha estimates through both artifact attenuation and removal of neural activity, and its performance depends on data length, channel density, decomposition quality, and classification thresholds. Moreover, the combination of ICA with absolute power was identified after inspection of the multiverse and requires independent validation. Other preprocessing choices can also influence EEG inference. Filtering and automatic rejection may alter the number of significant channels (Delorme, 2023), while reference performance depends on electrode coverage and density (Yao et al., 2019; Qin et al., 2010). Within the present decision space, however, filtering, ASR, and reference had smaller effects than ICA.

### 4.4. Spectral multiverse: absolute, relative, and aperiodic-adjusted power

Spectral estimation was the strongest post-preprocessing determinant of the *p*-value distribution. EEG spectra contain both oscillatory peaks and broadband aperiodic activity (Gyurkovics et al., 2021), which may arise from partly distinct physiological processes (Cross et al., 2025; Brake et al., 2024). Interpretation therefore depends on whether the spectrum is treated using additive or multiplicative models and whether the aperiodic component is retained or removed (Brake et al., 2024; Gyurkovics et al., 2021).

FOOOF and related methods separate periodic from aperiodic activity, which may improve mechanistic interpretation but not necessarily clinical discrimination. Aperiodic activity may itself contain disease-related information, and removing it could weaken a clinically useful marker of global slowing. Relative power may similarly reduce scaling differences while obscuring widespread spectral shifts when both numerator and denominator change. This may explain why absolute FFT and Welch estimates produced more stable clinical effects than relative or aperiodic-adjusted estimates. Accordingly, absolute power may maximize discriminability at the cost of mechanistic specificity.

Alpha definition, spatial region, and normalization can also influence EEG reliability and clinical discrimination (Chang, 2025; Bazanova, 2012). Standard terminology and frequency-band recommendations have therefore been proposed (Kane et al., 2017). However, fixed frequency boundaries may be less appropriate in neurodegenerative conditions characterized by slowing of the dominant rhythm. Such slowing has been documented in Parkinson’s and Alzheimer’s disease (Dauwels et al., 2010; Soikkeli et al., 1991), suggesting that individualized peak-frequency approaches may provide a useful extension. Within the present multiverse, however, changing the alpha upper boundary from 12 to 13 Hz or changing among Fz, Cz, frontal, and central summaries had less influence than spectral algorithm and power normalization.

### 4.5. Clinical Implications

The clinical value of the theta/alpha ratio depends not only on its association with cognitive impairment but also on whether that association survives reasonable analytic variation. In both datasets, higher theta/alpha ratios were most consistently associated with more advanced cognitive impairment, suggesting potential value as a severity-related qEEG marker rather than a diagnosis-specific test. However, the marked specification dependence means that a theta/alpha value cannot be interpreted clinically without considering how the EEG was processed. In particular, ICA combined with absolute FFT or Welch power emerged as a promising exploratory family, but these pipelines require prospective validation before they can support individual-level clinical decisions.

Multiverse analysis therefore has a practical role in biomarker development: it can identify which conclusions remain stable across defensible pipelines and which depend on narrow methodological choices. This is important for translating qEEG across laboratories, recording systems, and clinical settings, where differences in preprocessing can otherwise produce apparently conflicting results. Rather than selecting the pipeline with the strongest significance, candidate processing rules should be defined from multiverse results and tested prospectively in independent samples. Guided approaches may further reduce computational burden when the decision space becomes large (Dafflon et al., 2022). Such a workflow could help distinguish clinically transportable EEG markers from effects that are specific to one analytic implementation.

### 4.6. Limitations

Several limitations should be considered. First, Dataset 1 contained approximately 2 min of eyes-open EEG, whereas Dataset 2 contained approximately 12 to 14 min of eyes-closed EEG. Because eye state affects alpha activity and recording duration affects spectral stability, the datasets are not direct replications.

Second, sex distributions differed significantly across groups in both datasets. Because sex may influence EEG spectral measures, residual confounding may remain despite covariate adjustment, and generalizability should be interpreted cautiously. Adjusted and unadjusted models were both included for sensitivity analysis, although they are not necessarily equally valid when demographic imbalance is present.

Third, REST referencing should be interpreted cautiously in Dataset 2 because the 19-channel montage provides relatively sparse spatial sampling for a head-model-based reference.

Reference effects should therefore be viewed as specification sensitivity rather than evidence that REST performs equally across montages.

Fourth, the stronger performance of absolute FFT and Welch power relative to aperiodic-adjusted estimates may indicate that the theta/alpha ratio partly reflects broadband 1/f variation rather than purely oscillatory change. Absolute power may therefore improve discriminability while reducing mechanistic specificity.

Fifth, the 95% robustness threshold was descriptive rather than a formal statistical standard, and universes were dependent because they were derived from the same data. Proportions of significant universes should therefore be interpreted alongside effect direction and uncertainty.

Sixth, favorable combinations such as ICA with absolute FFT or Welch power were identified after inspection of the multiverse and require prospective validation. The multiverse also focused mainly on method-level choices and held many within-method parameters constant. Parameters such as filter order, ASR thresholds, ICA classification thresholds, spectral- window settings, and FOOOF parameters may affect results, but processing steps are interdependent, making exhaustive independent variation difficult and potentially generating implausible pipelines. The present findings therefore establish robustness only within the bounded decision space evaluated here.

Finally, the datasets differed in diagnosis, medication status, electrode density, and cognitive assessment, and PD cognitive groups were defined using MoCA thresholds rather than comprehensive clinical diagnostic procedures. These factors limit direct cross-dataset comparison.

## 5. Conclusions

This study used an exhaustive multiverse analysis to evaluate the EEG theta/alpha ratio across two neurodegenerative datasets. Pipeline families meeting the ≥95% robustness criterion also showed consistent effect directions, with higher theta/alpha ratios associated with greater cognitive impairment and lower cognitive scores. However, whether this criterion was reached depended strongly on analytical choices, particularly ICA and the use of absolute FFT- or Welch-derived power.

The theta/alpha ratio should therefore be considered a promising but specification-dependent marker rather than a pipeline-invariant biomarker. More broadly, the findings show how multiverse analysis can improve transparency in clinical EEG research by identifying both robust effects and the methodological conditions under which they emerge.

## Acknowledgements

None.

## CRediT author statement

**J.C.** conducted Conceptualization, Methodology, Software, Validation, Formal analysis, Investigation, Resources, Data Curation, Writing - Original Draft, Visualization, and Project administration. **S.H.S.** conducted Supervision and Writing - Review & Editing. All authors read and approved the final manuscript.

## Data Availability Statement

First dataset can be found in OpenNeuro: doi:10.18112/openneuro.ds004584.v1.0.0

Second dataset can be found in OpenNeuro: doi:10.18112/openneuro.ds004504.v1.0.9

## Abbreviations

AD: Alzheimer’s disease
ANCOVA: analysis of covariance
ANOVA: analysis of variance
ASR: Artifact Subspace Reconstruction
CTL: healthy controls
EEG: electroencephalography
FIR: finite impulse response
FOOOF: Fitting Oscillations and One-Over-F
FFT: fast Fourier transform
FTD: frontotemporal dementia
ICA: independent component analysis
LEDD: levodopa-equivalent daily dose
MMSE: Mini-Mental State Examination
MoCA: Montreal Cognitive Assessment
PD: Parkinson’s disease
PD-CN: cognitively normal Parkinson’s disease
PDD: Parkinson’s disease dementia
PDMCI: Parkinson’s disease with mild cognitive impairment
qEEG: quantitative electroencephalography
REST: reference electrode standardization technique
UPDRS-III: motor Unified Parkinson’s Disease Rating Scale.

## References

1. Romero, F. (2019). Philosophy of science and the replicability crisis. Philosophy Compass, 14(11).

2. Clayson, P. E. (2024). Beyond single paradigms, pipelines, and outcomes: Embracing multiverse analyses in psychophysiology. International Journal of Psychophysiology, 197, 112311–112311.

3. Delorme, A., & Makeig, S. (2004). EEGLAB: an open source toolbox for analysis of single-trial EEG dynamics including independent component analysis. Journal of Neuroscience Methods, 134(1), 9–21.

4. Schmidt, M. T., Kanda, P. A. M., Basile, L. F. H., da Silva Lopes, H. F., Baratho, R., Demario, J. L. C., Jorge, M. S., Nardi, A. E., Machado, S., Ianof, J. N., Nitrini, R., & Anghinah, R. (2013). Index of Alpha/Theta Ratio of the Electroencephalogram: A New Marker for Alzheimer’s Disease. Frontiers in Aging Neuroscience, 5.

5. Fahimi, G., Tabatabaei, S. M., Fahimi, E., & Rajebi, H. (2017). Index of theta/alpha ratio of the quantitative electroencephalogram in Alzheimer’s disease: a case-control study. Acta Medica Iranica, 502–506.

6. [dataset] Singh, A., Cole, R. C., Espinoza, A. I., Wessel, J. R., Cavanagh, J. F., & Narayanan, N. S. (2023). EEG: Parkinson’s disease, resting state (OpenNeuro dataset ds004584, version 1.0.0). OpenNeuro. 10.18112/openneuro.ds004584.v1.0.0

7. Kołodziej, A., Magnuski, M., Ruban, A., & Brzezicka, A. (2021). No relationship between frontal alpha asymmetry and depressive disorders in a multiverse analysis of five studies. Elife, 10, e60595.

8. Babiloni, C., Arakaki, X., Azami, H., Bennys, K., Blinowska, K., Bonanni, L., … & Guntekin, B. (2021). Measures of resting state EEG rhythms for clinical trials in Alzheimer’s disease: recommendations of an expert panel. Alzheimer’s & Dementia, 17(9), 1528–1553.

9. Dalrymple-Alford, J. C., MacAskill, M. R., Nakas, C. T., Livingston, L., Graham, C., Crucian, G. P., … & Anderson, T. J. (2010). The MoCA: well-suited screen for cognitive impairment in Parkinson disease. Neurology, 75(19), 1717–1725.

10. Goetz, C. G., Tilley, B. C., Shaftman, S. R., Stebbins, G. T., Fahn, S., Martinez Martin, P., … & LaPelle, N. (2008). Movement Disorder Society sponsored revision of the Unified Parkinson’s Disease Rating Scale (MDS UPDRS): scale presentation and clinimetric testing results. Movement disorders: official journal of the Movement Disorder Society, 23(15), 2129–2170.

11. Yao, D. (2001). A method to standardize a reference of scalp EEG recordings to a point at infinity. Physiological measurement, 22(4), 693–711.

12. [dataset] Miltiadous, A., Tzimourta, K. D., Afrantou, T., Ioannidis, P., Grigoriadis, N., Tsalikakis, D. G., Angelidis, P., et al. (2023). A dataset of EEG recordings from Alzheimer’s disease, frontotemporal dementia and healthy subjects (OpenNeuro dataset ds004504, version 1.0.9). OpenNeuro. 10.18112/openneuro.ds004504.v1.0.9

13. Kurlowicz, L., & Wallace, M. (1999). The Mini-Mental State Examination (MMSE). Journal of Gerontological Nursing, 25(5), 8–9.

14. Bell, C. C. (1994). DSM-IV: Diagnostic and statistical manual of mental disorders. JAMA, 272(10), 828–829.

15. McKhann, G., Drachman, D., Folstein, M., Katzman, R., Price, D., & Stadlan, E. M. (1984). Clinical diagnosis of Alzheimer’s disease: Report of the NINCDS-ADRDA Work Group under the auspices of Department of Health and Human Services Task Force on Alzheimer’s Disease. Neurology, 34(7), 939–944.

16. Botvinik-Nezer, R., Holzmeister, F., Camerer, C. F., Dreber, A., Huber, J., Johannesson, M., … & Rieck, J. R. (2020). Variability in the analysis of a single neuroimaging dataset by many teams. Nature, 582(7810), 84–88.

17. Vigário, R., Sarela, J., Jousmiki, V., Hamalainen, M., & Oja, E. (2000). Independent component approach to the analysis of EEG and MEG recordings. IEEE transactions on biomedical engineering, 47(5), 589–593.

18. Grech, R., Cassar, T., Muscat, J., Camilleri, K. P., Fabri, S. G., Zervakis, M., … & Vanrumste, B. (2008). Review on solving the inverse problem in EEG source analysis. Journal of neuroengineering and rehabilitation, 5(1), 25.

19. Romero, S., Mananas, M. A., Clos, S., Gimenez, S., & Barbanoj, M. J. (2003, September). Reduction of EEG artifacts by ICA in different sleep stages. In Proceedings of the 25th Annual International Conference of the IEEE Engineering in Medicine and Biology Society (EMBS’03) (Vol. 3, pp. 2675–2678).

20. Delorme, A. (2023). EEG is better left alone. Scientific reports, 13(1), 2372.

21. Yao, D., Qin, Y., Hu, S., Dong, L., Bringas Vega, M. L., & Valdés Sosa, P. A. (2019). Which reference should we use for EEG and ERP practice?. Brain topography, 32(4), 530–549.

22. Qin, Y., Xu, P., & Yao, D. (2010). A comparative study of different references for EEG default mode network: the use of the infinity reference. Clinical neurophysiology, 121(12), 1981–1991.

23. Gyurkovics, M., Clements, G. M., Low, K. A., Fabiani, M., & Gratton, G. (2021). The impact of 1/f activity and baseline correction on the results and interpretation of time-frequency analyses of EEG/MEG data: A cautionary tale. NeuroImage, 237, 118192.

24. Cross, Z. R., Gray, S. M., Dede, A. J., Rivera, Y. M., Yin, Q., Vahidi, P., … & Johnson, E. L. (2025). The development of aperiodic neural activity in the human brain. Nature human behaviour, 1–16.

25. Brake, N., Duc, F., Rokos, A., Arseneau, F., Shahiri, S., Khadra, A., & Plourde, G. (2024). A neurophysiological basis for aperiodic EEG and the background spectral trend. Nature communications, 15(1), 1514.

26. Donoghue, T., Haller, M., Peterson, E. J., Varma, P., Sebastian, P., Gao, R., … & Voytek, B. (2020). Parameterizing neural power spectra into periodic and aperiodic components. Nature neuroscience, 23(12), 1655–1665.

27. Dafflon, J., Da Costa, P. F., Váša, F., Monti, R. P., Bzdok, D., Hellyer, P. J., … & Leech, R. (2022). A guided multiverse study of neuroimaging analyses. Nature Communications, 13(1), 3758.

28. Chang, J. (2025). Test Retest Reliability of Single Spectral Power and Spectral Power Ratios in Relative and Absolute Values. Brain and Behavior, 15(11), e71035.

29. Bazanova, O. (2012). Comments for current interpretation EEG alpha activity: a review and analysis. Journal of Behavioral and Brain Science, 2(2), 239–248.

30. Kane, N., Acharya, J., Beniczky, S., Caboclo, L., Finnigan, S., Kaplan, P. W., … & Van Putten, M. J. (2017). A revised glossary of terms most commonly used by clinical electroencephalographers and updated proposal for the report format of the EEG findings. Revision 2017. Clinical neurophysiology practice, 2, 170–185.

31. Dauwels, J., Vialatte, F., & Cichocki, A. (2010). Diagnosis of Alzheimer’s disease from EEG signals: where are we standing?. Current Alzheimer Research, 7(6), 487–505.

32. Soikkeli, R., Partanen, J., Soininen, H., Pääkkönen, A., & Riekkinen Sr, P. (1991). Slowing of EEG in Parkinson’s disease. Electroencephalography and clinical neurophysiology, 79(3), 159–165.

33. Liang, Y., Li, P., Wang, Y., Jing, F., Dong, C., & Zhao, L. (2026). Application of resting-state EEG theta/alpha power ratio analysis for diagnosing amnestic mild cognitive impairment. Scientific Reports.

34. Hamilton, C. A., Schumacher, J., Matthews, F., Taylor, J. P., Allan, L., Barnett, N., … & Thomas, A. J. (2021). Slowing on quantitative EEG is associated with transition to dementia in mild cognitive impairment. International Psychogeriatrics, 33(12), 1321–1325.

35. Yao, D., Wang, L., Oostenveld, R., Nielsen, K. D., Arendt-Nielsen, L., & Chen, A. C. (2005). A comparative study of different references for EEG spectral mapping: the issue of the neutral reference and the use of the infinity reference. Physiological measurement, 26(3), 173–184.

36. Gibb, W. R., & Lees, A. (1988). The relevance of the Lewy body to the pathogenesis of idiopathic Parkinson’s disease. *Journal of Neurology*, Neurosurgery & Psychiatry, 51(6), 745–752.

37. Widmann, A., Schröger, E., & Maess, B. (2015). Digital filter design for electrophysiological data–a practical approach. Journal of neuroscience methods, 250, 34–46.

38. Winkler, I., Debener, S., Müller, K. R., & Tangermann, M. (2015, August). On the influence of high-pass filtering on ICA-based artifact reduction in EEG-ERP. In 2015 37th annual international conference of the IEEE engineering in medicine and biology society (EMBC) (pp. 4101–4105). IEEE.

39. Chang, C. Y., Hsu, S. H., Pion-Tonachini, L., & Jung, T. P. (2019). Evaluation of artifact subspace reconstruction for automatic artifact components removal in multi-channel EEG recordings. IEEE transactions on biomedical engineering, 67(4), 1114–1121.

40. Jensen, O., & Tesche, C. D. (2002). Frontal theta activity in humans increases with memory load in a working memory task. European journal of Neuroscience, 15(8), 1395–1399.

41. Cavanagh, J. F., & Frank, M. J. (2014). Frontal theta as a mechanism for cognitive control. Trends in cognitive sciences, 18(8), 414–421.

42. Allen, J. J., Keune, P. M., Schönenberg, M., & Nusslock, R. (2018). Frontal EEG alpha asymmetry and emotion: From neural underpinnings and methodological considerations to psychopathology and social cognition. Psychophysiology, 55(1), e13028.

43. Singh, A., Richardson, S. P., Narayanan, N., & Cavanagh, J. F. (2018). Mid-frontal theta activity is diminished during cognitive control in Parkinson’s disease. Neuropsychologia, 117, 113–122.

44. Robbins, K. A., Touryan, J., Mullen, T., Kothe, C., & Bigdely-Shamlo, N. (2020). How sensitive are EEG results to preprocessing methods: a benchmarking study. IEEE transactions on neural systems and rehabilitation engineering, 28(5), 1081–1090.

45. Huang, H., Moffa, A. H., Loo, C., & Nikolin, S. (2025). No Single Best Pipeline: Multiverse Analysis of EEG Preprocessing for N Back Working Memory Tasks. Psychophysiology, 62(12), e70197.

46. Wiesman, A. I., da Silva Castanheira, J., & Baillet, S. (2022). Stability of spectral estimates in resting-state magnetoencephalography: Recommendations for minimal data duration with neuroanatomical specificity. Neuroimage, 247, 118823.

47. Miyakoshi, M. (2023). Artifact subspace reconstruction: a candidate for a dream solution for EEG studies, sleep or awake. Sleep, 46(12), zsad241.

48. İşoğlu-Alkaç, Ü., & Strüber, D. (2006). Necker cube reversals during long-term EEG recordings: sub-bands of alpha activity. International Journal of Psychophysiology, 59(2), 179–189.

49. Kopčanová, M., Tait, L., Donoghue, T., Stothart, G., Smith, L., Flores-Sandoval, A. A., … & Benwell, C. S. (2024). Resting-state EEG signatures of Alzheimer’s disease are driven by periodic but not aperiodic changes. Neurobiology of Disease, 190, 106380.

50. Gudmundsson, S., Runarsson, T. P., Sigurdsson, S., Eiriksdottir, G., & Johnsen, K. (2007). Reliability of quantitative EEG features. Clinical Neurophysiology, 118(10), 2162–2171.

