## Supplementary Figures for "Specification dependence of EEG theta/alpha ratio associations with cognitive impairment: a multiverse analysis"

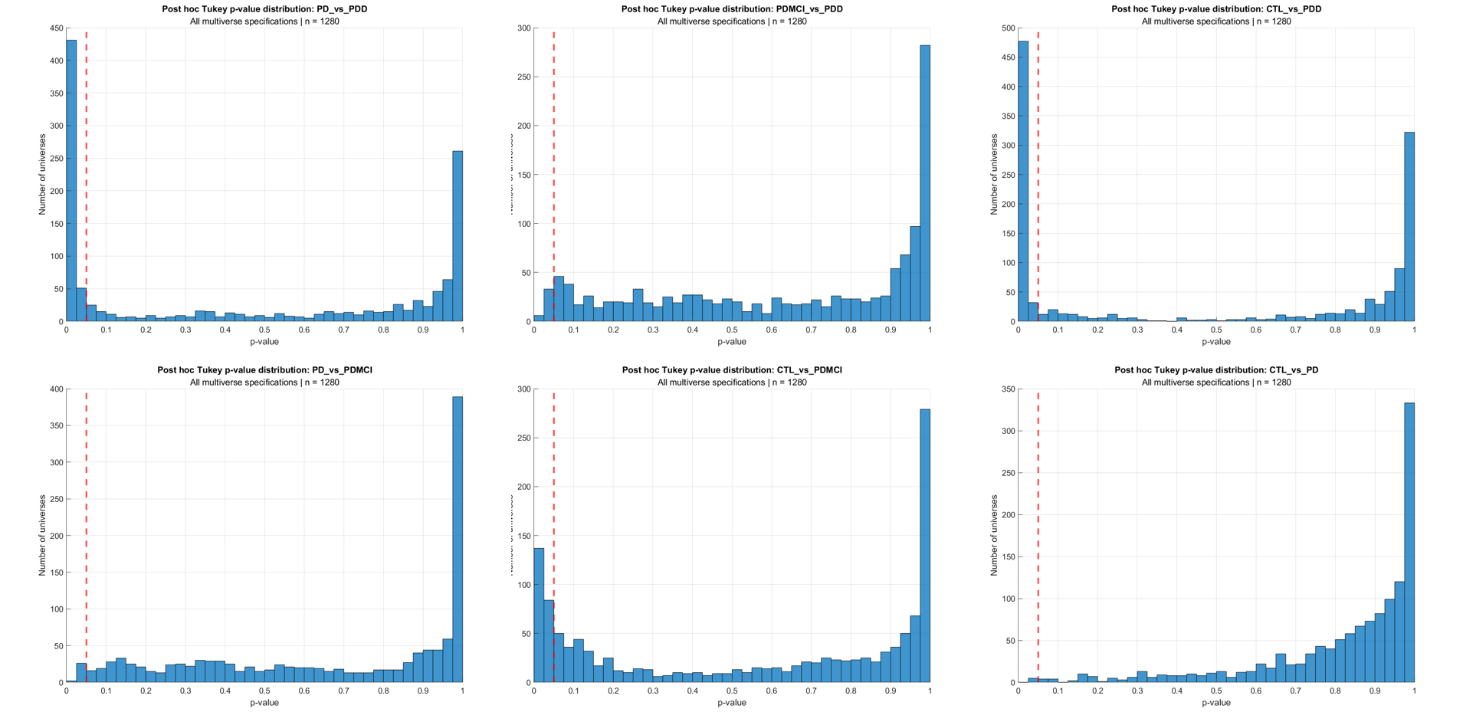

**Supplementary Figure 1. Distribution of pairwise p values for theta/alpha ratio comparisons among CTL, PD-CN (PD), PDMCI, and PDD groups across the full multiverse.** The vertical red line indicates p = 0.05.

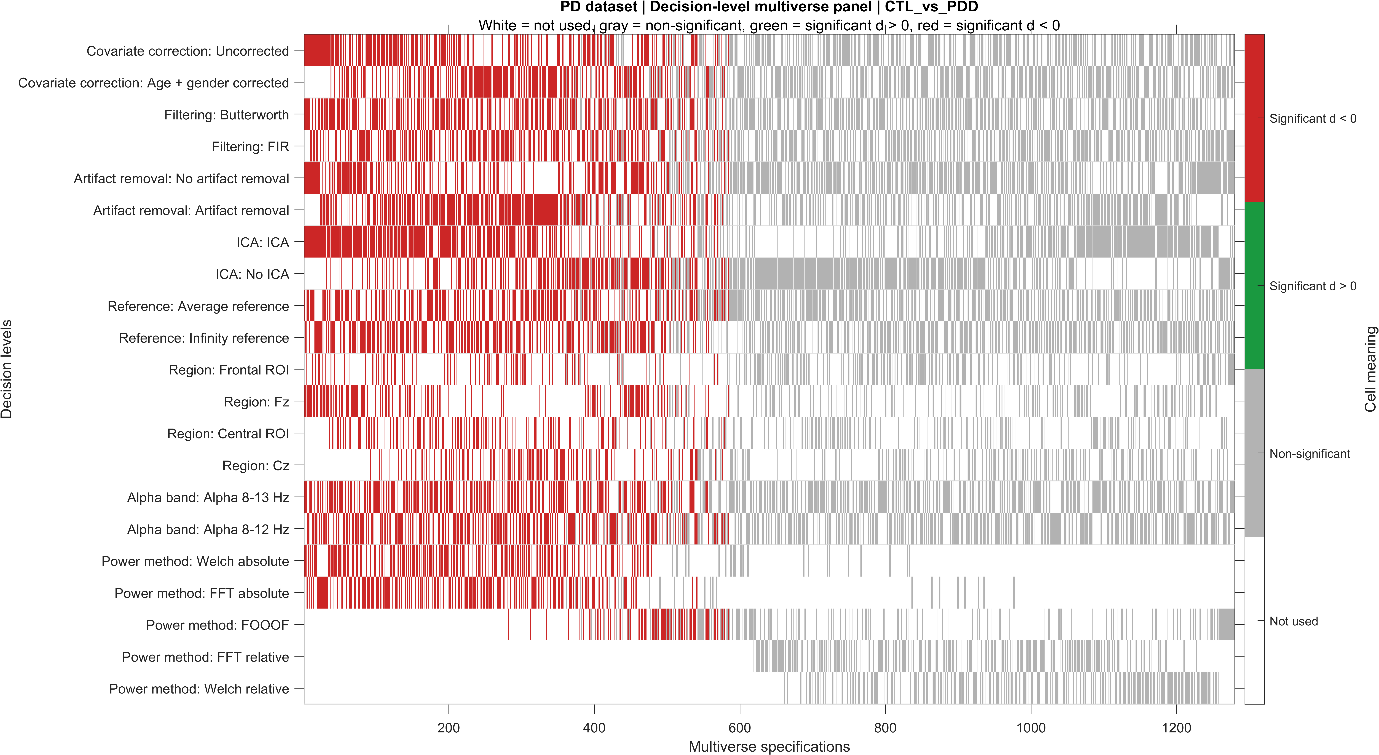

**Supplementary Figure 2. Specification curve of all universes for the CTL versus PDD comparison stratified by each analytic decision.** Cohen’s d was estimated as effect sizes over each decision point. Statistically positive estimate was marked as green (CTL > PDD), while statistically negative estimate was marked as red (CTL < PDD) and gray represents nonsignificant.

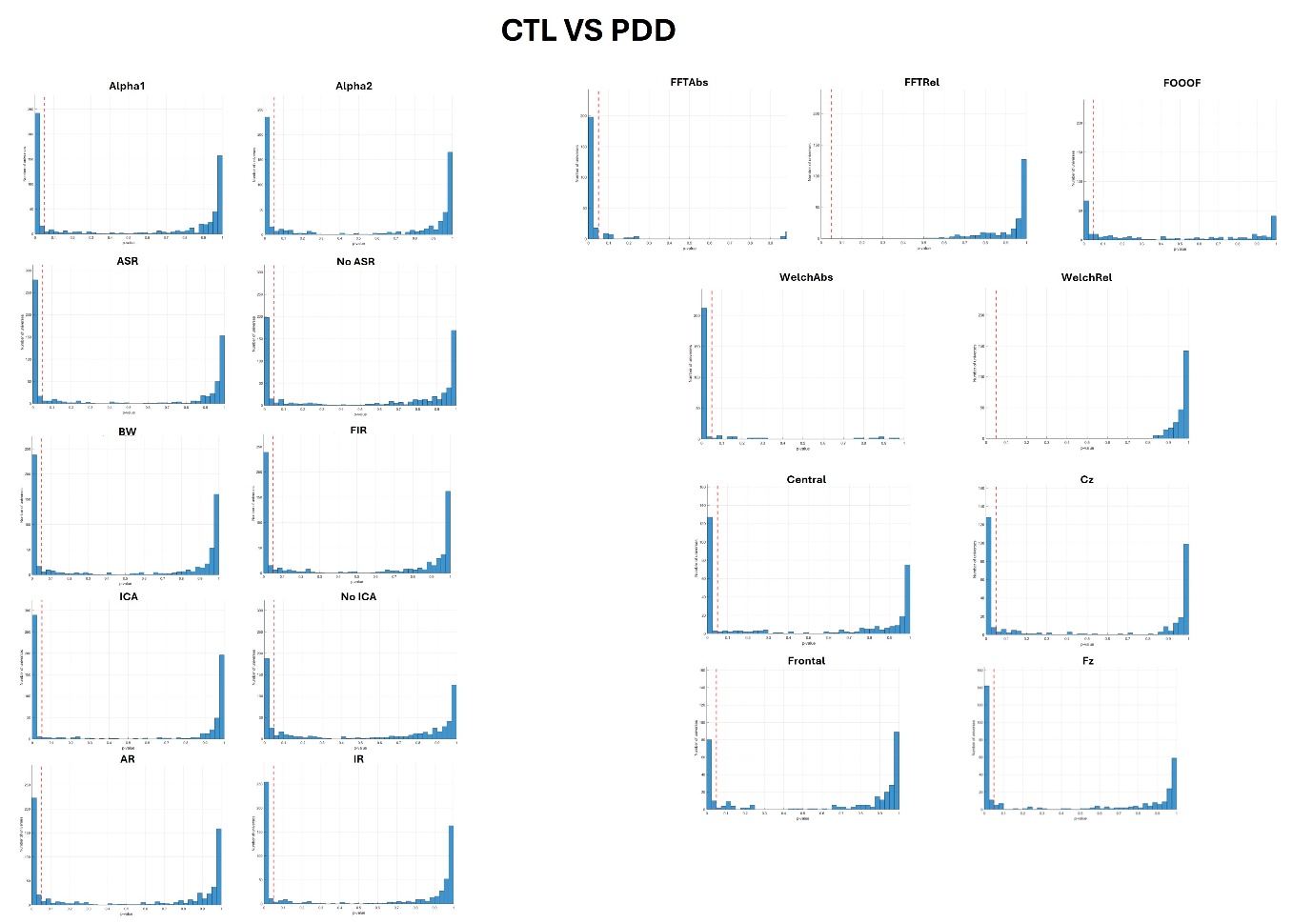

**Supplementary Figure 3. Distribution of p values for the CTL versus PDD comparison stratified by each analytic decision.** The vertical red line indicates p = 0.05. Alpha 1: 8-12Hz; Alpha 2: 8-13Hz; ASR: Artifact Subspace Reconstruction; BW: Butterworth; FIR: Finite Impulse Response; ICA: Independent Component Analysis; AR: Average Reference; IR: Infinity Reference; Abs: absolute power; Rel: relative power;

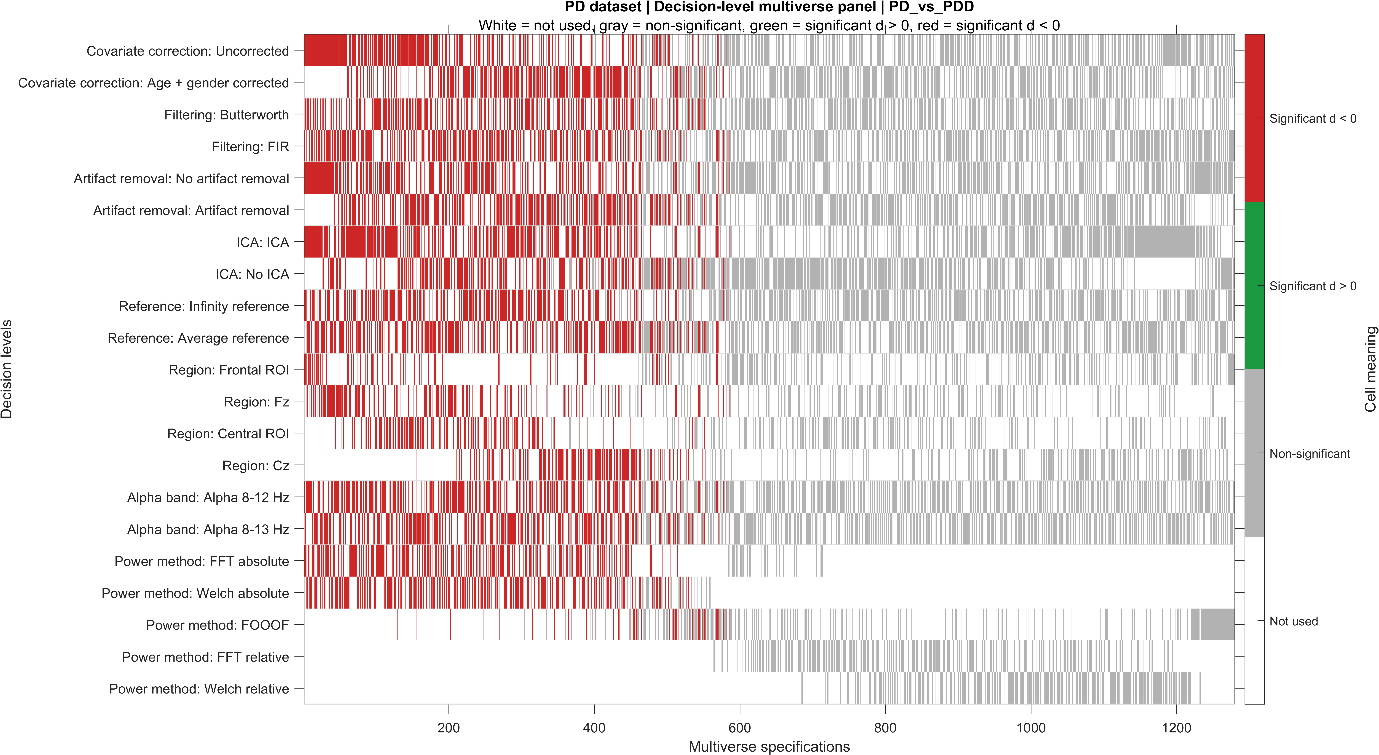

**Supplementary Figure 4. Specification curve of all universes for the PD-CN (PD) versus PDD comparison stratified by each analytic decision.** Cohen’s d was estimated as effect sizes over each decision point. Statistically positive estimate was marked as green (PD > PDD), while statistically negative estimate was marked as red (PD < PDD) and gray represents nonsignificant.

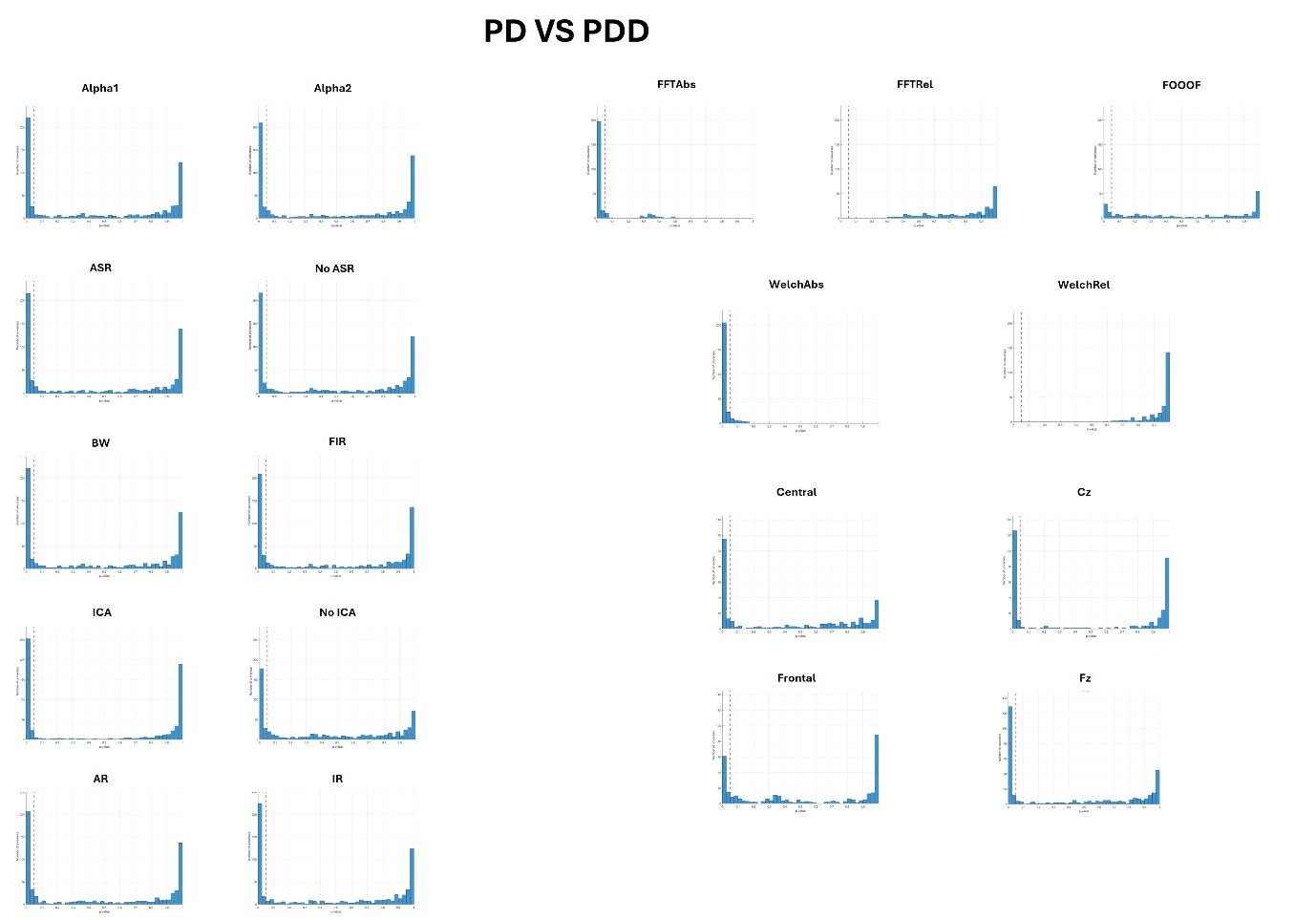

**Supplementary Figure 5. Distribution of p values for the PD-CN (PD) versus PDD comparison stratified by each analytic decision.** The vertical red line indicates p = 0.05. Alpha 1: 8-12Hz; Alpha 2: 8-13Hz; ASR: Artifact Subspace Reconstruction; BW: Butterworth; FIR: Finite Impulse Response; ICA: Independent Component Analysis; AR: Average Reference; IR: Infinity Reference; Abs: absolute power; Rel: relative power;

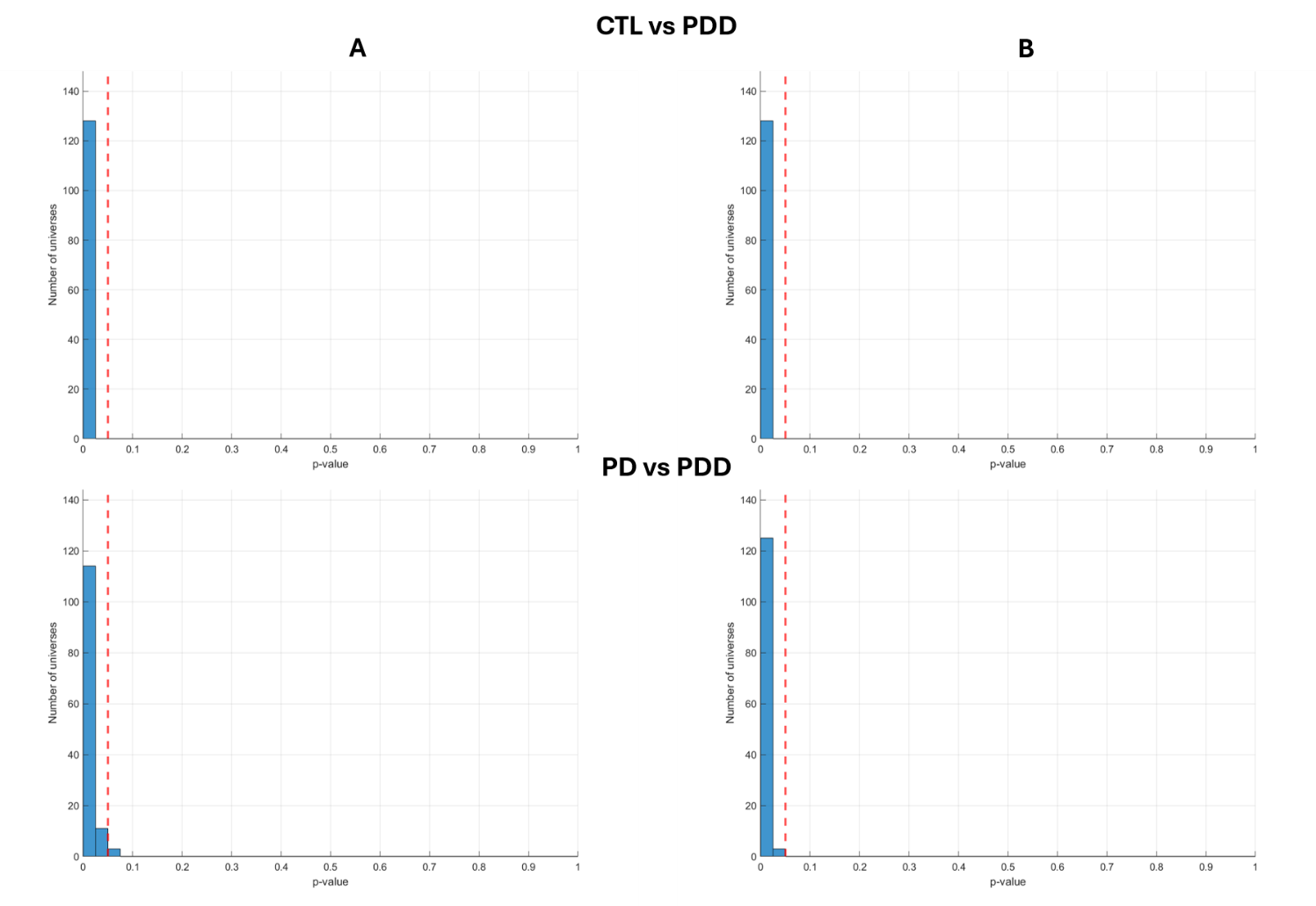

**Supplementary Figure 6. Distribution of p values for the CTL versus PDD and PD-CN (PD) versus PDD comparison among universes that included ICA and used (A) absolute FFT power or (B) absolute Welch power.** The vertical red line indicates p = 0.05.

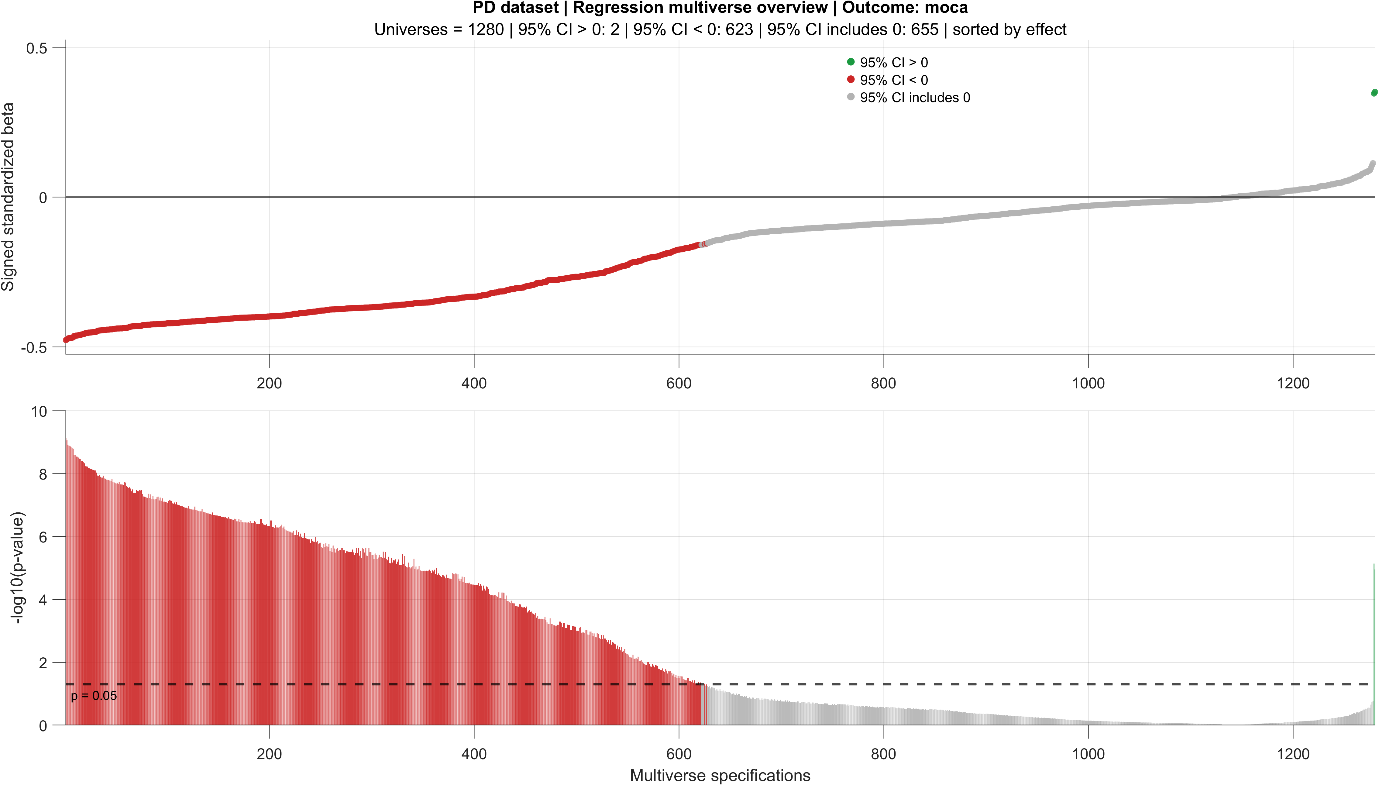

**Supplementary Figure 7. Distribution of effect sizes and p values from linear regression analyses relating the theta/alpha ratio to MoCA scores across the full multiverse.** Red represents the negative relationship.

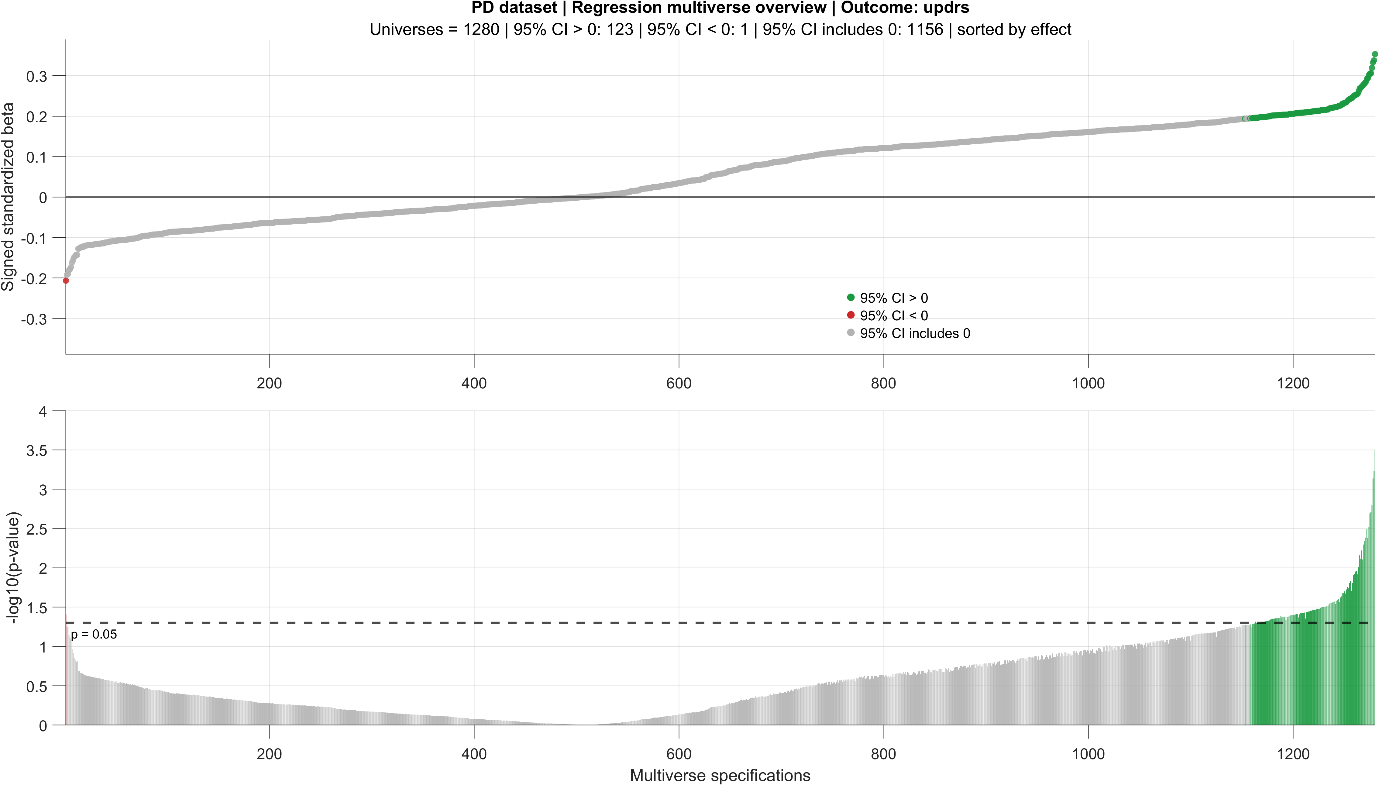

**Supplementary Figure 8. Distribution of effect sizes and p values from linear regression analyses relating the theta/alpha ratio to UPDRS-III scores across the full multiverse.** Green represents the positive relationship.

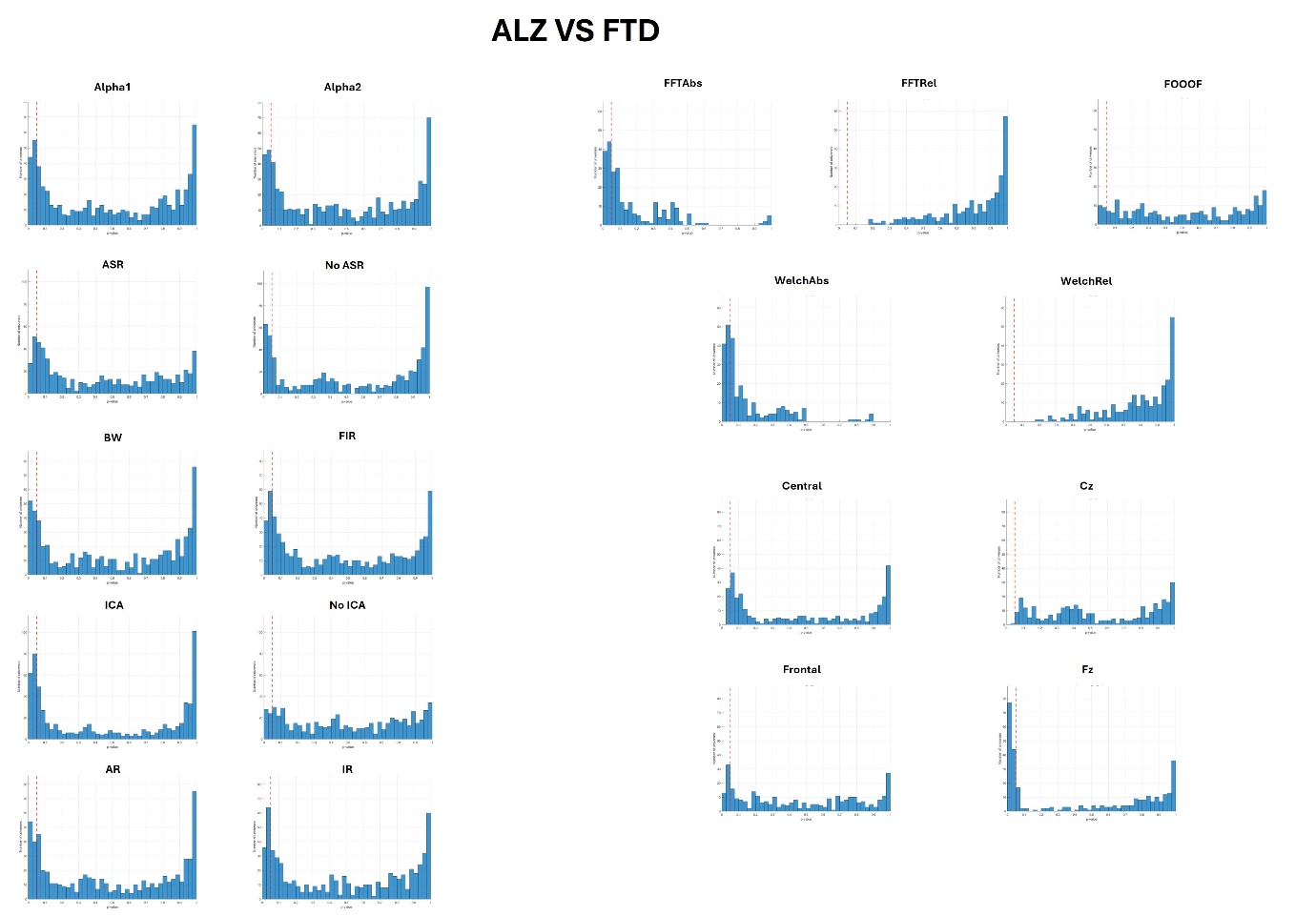

**Supplementary Figure 9. Distribution of p values for the AD versus FTD comparison stratified by each analytic decision.** The vertical red line indicates p = 0.05. Alpha 1: 8-12Hz; Alpha 2: 8-13Hz; ASR: Artifact Subspace Reconstruction; BW: Butterworth; FIR: Finite Impulse Response; ICA: Independent Component Analysis; AR: Average Reference; IR: Infinity Reference; Abs: absolute power; Rel: relative power;

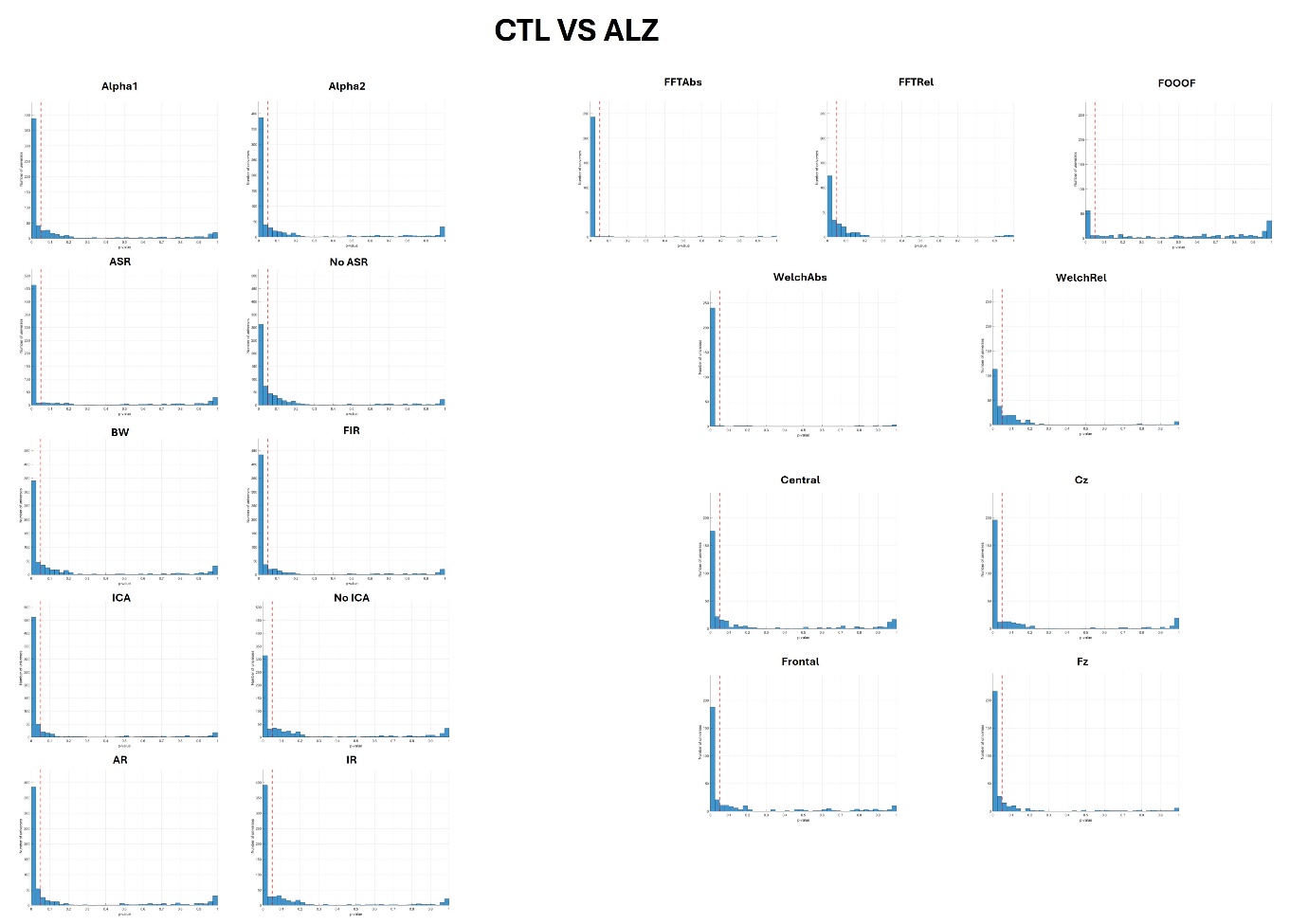

**Supplementary Figure 10. Distribution of p values for the CTL versus AD comparison stratified by each analytic decision.** The vertical red line indicates p = 0.05. Alpha 1: 8-12Hz; Alpha 2: 8-13Hz; ASR: Artifact Subspace Reconstruction; BW: Butterworth; FIR: Finite Impulse Response; ICA: Independent Component Analysis; AR: Average Reference; IR: Infinity Reference; Abs: absolute power; Rel: relative power;

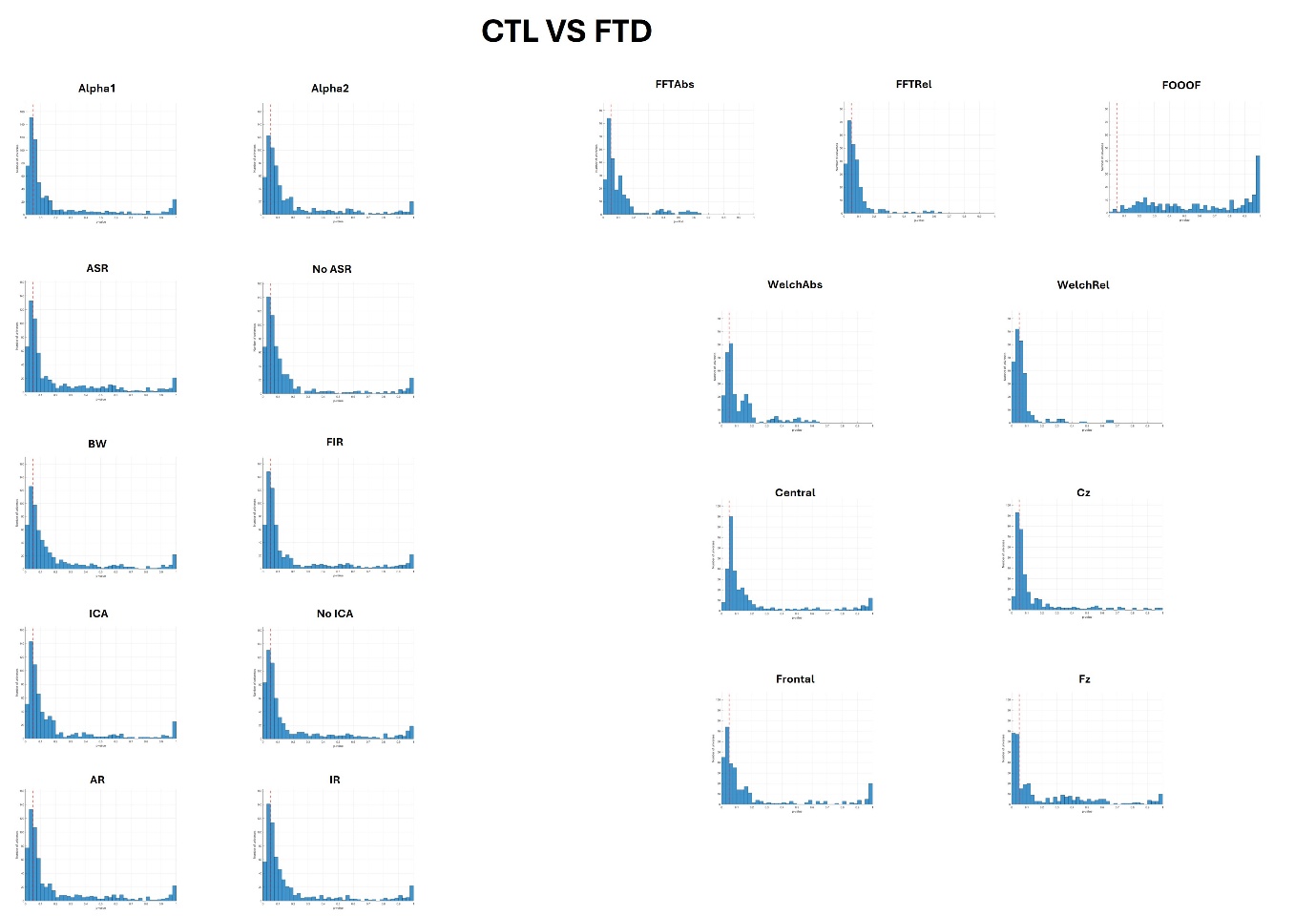

**Supplementary Figure 11. Distribution of p values for the CTL versus FTD comparison stratified by each analytic decision.** The vertical red line indicates p = 0.05. Alpha 1: 8-12Hz; Alpha 2: 8-13Hz; ASR: Artifact Subspace Reconstruction; BW: Butterworth; FIR: Finite Impulse Response; ICA: Independent Component Analysis; AR: Average Reference; IR: Infinity Reference; Abs: absolute power; Rel: relative power;
